# Small non-coding RNA landscape of extracellular vesicles from a post-traumatic model of equine osteoarthritis

**DOI:** 10.1101/2022.03.10.483752

**Authors:** James R Anderson, Stine Jacobsen, Marie Walters, Louise Bundgaard, Andreas Diendorfer, Matthias Hackl, Emily J Clarke, Victoria James, Mandy J Peffers

**Affiliations:** Musculoskeletal Ageing Science, Institute of Life Course and Medical Sciences, University of Liverpool, UK; Department of Veterinary Clinical Sciences, University of Copenhagen, Taastrup, Denmark; Tamirna, TAmiRNA GmbH, Vienna, Austria; University of Nottingham

## Abstract

Extracellular vesicles comprise an as yet inadequately investigated intercellular communication pathway in the field of early osteoarthritis. We hypothesised that small non-coding RNA expression pattern in synovial fluid and plasma would change during progression of experimental osteoarthritis. In this study, we used small RNA sequencing to provide a comprehensive overview of the temporal expression profiles of small non-coding transcripts carried by EVs derived from plasma and synovial fluid for the first time in a post-traumatic model of equine osteoarthritis. Additionally, we characterised synovial fluid and plasma-derived extracellular vesicles with respect to quantity, size, and surface markers. The differential expression of seven microRNAs in plasma and synovial fluid-derived extracellular vesicles; miR-451, miR-25, miR-215, miR-92a, miR-let-7c, miR-486-5p, miR-23a and four snoRNAs; U3, snord15, snord46, snord58 represent potential biomarkers for early OA. Bioinformatics analysis of the differentially expressed microRNAs in synovial fluid highlighted that in early OA these related to the inhibition of cell cycle, cell cycle progression, DNA damage and cell proliferation but increased cell viability, and differentiation of stem cells. Plasma and synovial fluid-derived extracellular vesicle small non-coding signatures have been established for the first time in a temporal model of osteoarthritis. These could serve as novel biomarkers for the evaluation of osteoarthritis progression or act as potential therapeutic targets.

## Introduction

Osteoarthritis (OA) is a degenerative joint disease characterised by deterioration of articular cartilage, accompanied by changes in the bone and soft tissues of the joint (1) which adversely impacts the health of the equine athlete. It is a major welfare issue resulting in substantial morbidity and mortality (2). Lameness resulting from OA is a major cause of poor performance and early retirement (3). Despite the huge socioeconomic importance of OA, our understanding of the pathophysiological mechanisms involved is limited (4). OA is characterised by an increase in cartilage extracellular matrix (ECM) degradation by proteases and a reduction in ECM production (5). We recently identified that differential expression (DE) of small nucleolar RNAs (snoRNAs) contributes to this imbalance; a key mechanism in OA (6, 7). We require biomarkers to identify early OA before cartilage ECM is irreversibly degraded and our group has identified small non-coding RNAs (sncRNAs) distinguishing early equine OA synovial fluid (SF) (8).

SncRNAs such as microRNAs (miRs) and snoRNAs are functional RNA molecules that are transcribed from DNA but do not translate into proteins (9). Furthermore, it is expected that novel therapies can be directed in a personalised manner (10) following disease stratification. This is imperative as OA therapies are currently only symptomatic; principally pain relief in the horse. Extracellular vesicles (EVs) from purified cell types are suggested as novel therapeutics in a number of human diseases including rheumatoid arthritis (11), cardiac disease (12) and neoplasia (13).

We have identified DE sncRNAs in ageing and OA cartilage (14, 15), OA SF (8) and serum (16, 17). In human OA others have identified SNCRNAs in plasma and SF as potential biomarkers (18–22), as well as plasma EVs (23). EVs produced by cells, transfer molecules (including SNCRNAs) between cells and tissues (24) and are found in serum, SF, articular cartilage and supernatants of synoviocytes and chondrocytes (25). EV cargo is involved in cross-talk between cells within joint tissues and affects ECM turnover and inflammation(26, 27), representing a crucial step in OA regulation. The role of EVs in OA provides a foundation to create novel disease-modifying treatments (26). Promising results were obtained in the therapeutic application of mesenchymal stem cell-derived EVs for cartilage repair and experimental OA (28). Additionally, EVs have therapeutic potential in rheumatoid arthritis when administered either into the joint or systemically (26).

Animal models of OA enable the reproduction and progression of degenerative damage to be measured in a controlled manner, enabling opportunities to monitor and modulate symptoms and disease progression (29). Although the equine carpal osteochondral fragment model (30, 31) does not encompass all pathophysiological aspects of different OA phenotypes, it allows a controlled system with known time of onset and a singular cause of OA, facilitating temporal OA progression studies. Molecules can be measured temporally to determine their role in early OA. This is particularly imperative as the precise onset of molecular events of early OA following trauma is not completely understood. Furthermore, the size of the equine middle carpal joint permits repeated SF sampling over time, thus limiting use of experimental animals.

We hypothesise that EV sncRNA cargo from SF and plasma can be used to identify OA at an early stage before clinical signs and irreversible cartilage degradation. Additionally, by determining changes in sncRNAs in a longitudinal manner we may be able to further understand the pathogenesis of early OA.

## Methods

### Horses and study design

The Danish Animal Experiments Inspectorate (permit 2017-15-0201-01314) and local ethical committee of the Large Animal Teaching Hospital of University of Copenhagen approved the experimental protocol. All procedures were undertaken according to EU Directive 2010/63/EU for animal experiments.

Four skeletally mature Standardbred trotters (2.5-7 years old, weighing 397-528 kg) were included in this study. Prior to inclusion, the horses underwent clinical examination, lameness examination including flexion tests, radiographic imaging, haematological and blood-biochemical analysis, and arthrocentesis of both middle carpal joints to ensure that horses were sound and joints healthy.

OA was surgically induced in the left middle carpal joint and the right middle carpal joint underwent sham surgery as described previously (31) and plasma plus SF; sampled from both middle carpals before and following OA induction as described below.

The horses were euthanised on day 71/72 with pentobarbital sodium (140 mg/kg, Euthasol Vet, Dechra Veterinary Products, Uldum, Denmark). Following euthanasia, samples were collected from the joints as detailed below.

### Induction of osteoarthritis and exercise

OA induction was undertaken as described by McIlwraith (31) under general anaesthetic. Sham surgery (arthroscopy alone) was performed in the right middle carpal joint. All portals were sutured using 2-0 propylene (Surgipro, Medtronic Danmark A/S, Copenhagen, Denmark). After 2 weeks following OA induction horses were exercised in 2 min of trot (4.4-5.3 m/sec), 2 min of fast trot/gallop (9 m/sec) and 2 min of trot (4.4-5.3 m/sec) for 5 days/week on a treadmill (32).

### SF and plasma sampling

SF samples were obtained from both middle carpal joints prior to (day 0) and 10, 35, 42, 49, 56, 63 after surgery. Following sedation with detomidine (0.01 mg/kg, Domosedan Vet, Orion Pharma Animal Health, Copenhagen, Denmark) and butorphanol tartrate (0.01 mg/kg, Dolorex, MSD Animal Health, Copenhagen, Denmark), SF was aspirated aseptically with a 19-gauge 40 mm needle and transferred to tubes containing ethylene diamine tetra acetic acid (EDTA) (BD Vacutainer, BD A/S, Albertslund, Denmark). These were inverted 5-10 times, stored on melting ice until centrifugation at 1000g for 20 min at 4 °C. Plasma was collected before sedation from the jugular vein. From day 0-10 samples were collected through an indwelling catheter where the first 10 mL were discharged and the sample transferred to a EDTA tube. For the rest of the study period samples were sampled directly into the EDTA tube using a vacutainer system. Tubes were inverted 5-10 times, centrifuged at 3000 x g for 15 min at 20 °C. Biofluids were processed within one hour and stored at −80 °C.

### Post-mortem examination

Following euthanasia, the middle carpal joints were opened and synovial membrane and articular cartilage obtained from the intermediate carpal and third carpal bone, and placed in neutral buffered 10 % formalin. For histology samples were processed for hematoxylin and eosin and safranin O (cartilage only) staining. Grading of the synovial membrane and cartilage was performed (33).

### EV Isolation

SF and plasma collected at day 0, 10, 35, 42, 49, 56, 63 with relevant controls were thawed and SF subsequently treated with 1 μg/ml hyaluronidase (from bovine testes, Sigma-Aldrich, Gillingham, UK). Both SF and plasma were centrifuged at 2,500*g* for 10 min and 10,000*g* for 10 min. EVs were subsequently isolated using size exclusion chromatography using qEVsingle columns (IZON, Lyon, France) following the manufacturer’s instructions. Briefly, 3.5 ml of phosphate buffered saline (PBS) (Sigma-Aldrich, Gillingham, UK), previously processed using a 0.22 μm polyethersulfone filter (Sartorius, Göttingen, Germany) was passed through the column, followed by 150 μl of SF or plasma. The first five 200 μl flow through fractions were discarded and the following five 200 μl fractions pooled (isolated EVs).

### EV Characterisation

#### Nanoparticle tracking analysis

For all samples, 100 μl of isolated EVs were characterised via nanoparticle tracking analysis (NTA) (Nanosight NS300, Malvern Panalytical Ltd, Malvern, UK) at 25°C to determine particle size and EV concentration. Samples were diluted 1:100 in non-sterile filtered PBS for the optimal particle concentration range of 107-109/ml (34). For each sample and control, three 60 s videos were taken with the sample advanced though the machine between each video. EVs were viewable with camera level 12 and screen gain 4, and the temperature was maintained at 25°C. Between each sample, the Nanosight chamber was flushed twice with 1 ml of PBS to prevent cross-contamination. Data analysis was performed with NTA 3.2 software (Malvern Panalytical, Malvern, UK).

#### Exoview

SF and plasma EV isolations collected at 0, 42 and 63 days were concentrated using 2 ml Vivaspin concentrator columns (Sartorius, Göttingen, Germany) by centrifuging at 1,000g until the final volume was 100 μl and subsequently centrifuged into the column cap at 4,000g for 2 min. 5 μl of each sample was removed and pooled with all corresponding samples of the same group and time point. ExoView analyses EVs using visible light interference for size measurements and fluorescence for protein profiling. Samples were analyzed in triplicate replicates /sample using the human ExoView Tetraspanin Kit (NanoView Biosciences, USA). Samples were diluted in manufacturer supplied incubation solution, and incubated overnight at room temperature on ExoView Tetraspanin Chips. Chips were washed three times in solution A, prior to incubation with fluorescent tetraspanin antibodies. Labelling antibodies consisted of anti-CD9 CF488, anti-CD81 CF555 and anti-CD63 CF647 and the MIgG negative control. Antibodies were diluted 1:500 as per manufacturer’s instructions and incubated on chips for 1 hour at room temperature. Chips were then washed in kit supplied buffers, dried and imaged by the ExoView R100 using ExoView Scanner v3.0. Data was analysed using ExoView Analyzer v3.0. Fluorescent cut offs were set relative to the MIgG control. Total EVs were determined as the number of detected particles bound to tetraspanin antibodies (CD9, CD81, CD63) and normalised to MIgG antibody. Particle diameter and counts were statistically analysed using repeated measures ANOVA in GraphPad Prism v9.0.1 and Excel. Significant differences between groups were identified with P ≤ 0.05.

### Small RNA sequencing

#### RNA isolation, library preparation for small RNA-Seq and sequencing

Total RNA was extracted from 200 μl of isolated EVs using the miRNeasy Serum/Plasma Advanced Kit (Qiagen, Crawley, UK) following the manufacturer’s instructions. Due to the low RNA yield, which impairs accurate determination of RNA concentrations, a fixed total RNA volume of 2 μl was used for library preparation using the CleanTag kit (TriLink, San Diego, USA). Adapter dilution (1:4) and PCR amplification (26 cycles) were optimized in a pilot experiment using Bioanalyzer DNA1000 (Agilent, Sant Clara, USA) chips as a read out of library size and quantity. Samples were then processed in two batches á 37 samples. Library yield and size range was confirmed for all samples using Bioanalyzer DNA1000 chips (Agilent, Santa Clara, USA). Equimolar amounts of all samples were pooled and size purification was performed using BluePippin (SageBiosystems, Beverly, USA). To maintain larger RNA fragments the size range was extend to 130bp – 300bp, which corresponds to insert sizes between 15bp and 180bp. Purified pool were sequenced on a NovaSeq SP100 flow cell (Illumina, San Diego, USA).

### Small RNA sequencing data processing

Data was analysed using the miND pipeline. Overall quality of the next-generation sequencing data was evaluated automatically and manually with fastQC v0.11.9 (35) and multiQC v1.10 (36). Reads from all passing samples were adapter trimmed and quality filtered using cutadapt v3.3 (37) and filtered for a minimum length of 17nt. Mapping steps were performed with bowtie v1.3.0 (38)and miRDeep2 v2.0.1.2 (39), whereas reads were mapped first against the genomic reference EquCab.3.0 provided by Ensembl (40) allowing for two mismatches and subsequently miRBase v22.1 (41), filtered for miRs of *Equus caballus* only, allowing for one mismatch. For a general RNA composition overview, non-miR mapped reads were mapped against RNAcentral (42) and then assigned to various RNA species of interest. Statistical analysis of pre-processed next generation sequencing data was undertaken with R v4.0 and the packages pheatmap, pcaMethods v1.82 and genefilter v1.72. DE analysis with edgeR v3.32 (43) used the quasi-likelihood negative binomial generalized log-linear model functions provided by the package. The independent filtering method of DESeq2 (44) was adapted for use with edgeR to remove low abundant miRs and thus optimize the false discovery rate (FDR) correction. Significantly DE sncRNAs were determined as P<0.05.

### Data availability

Raw NGS data is available at GEO ID #######.

### Bioinformatics of differentially expressed microRNAs

Potential biological associations of the DE miRs in EVs derived from plasma or SF were identified using Ingenuity Pathway Ananalysis (IPA) (IPA, Qiagen Redwood City, CA, USA) ‘Core Analysis’. Furthermore, to identify putative miR targets, we utilised MicroRNA Target Filter module within IPA. We used a conservative filter; experimentally validated and highly conserved predicted mRNA targets for each miRNA. ToppGene was used for functional enrichment analysis of the miRNA targets using ToppGene (45) with a Bonferroni FDR of less than 0.05. Biological process gene ontology (GO) terms generated through ToppGene were then summarised, and REViGO (46) and Cytoscape (47) used to visualise the network.

## Results

### Model outcome

As previously described the end point synovial membrane scores were significant increased, with increased cellular infiltration, intimal hyperplasia, and subintermal oedema in the OA joints versus control joints (p <0.05). Additionally previous histological evaluation of the third carpal bone cartilage demonstrated a significantly increase in chondrocyte necrosis, cluster formation, and focal cell loss scores in OA joints, with a significant increase in final score (p < 0.05) (48).

### EV characterisation

#### Nanoparticle tracking analysis

The mean and mode particle size for control SF was (219.8, 180.9nm), OA SF (221.8, 195nm) and plasma (158.4, 123.3nm). We did not identify differences for plasma or SF in size, size distribution or concentration between control and OA joints nor differences between the sampled time points (Figure 1, Supplementary Files 1 and 2).

**Figure 1.**
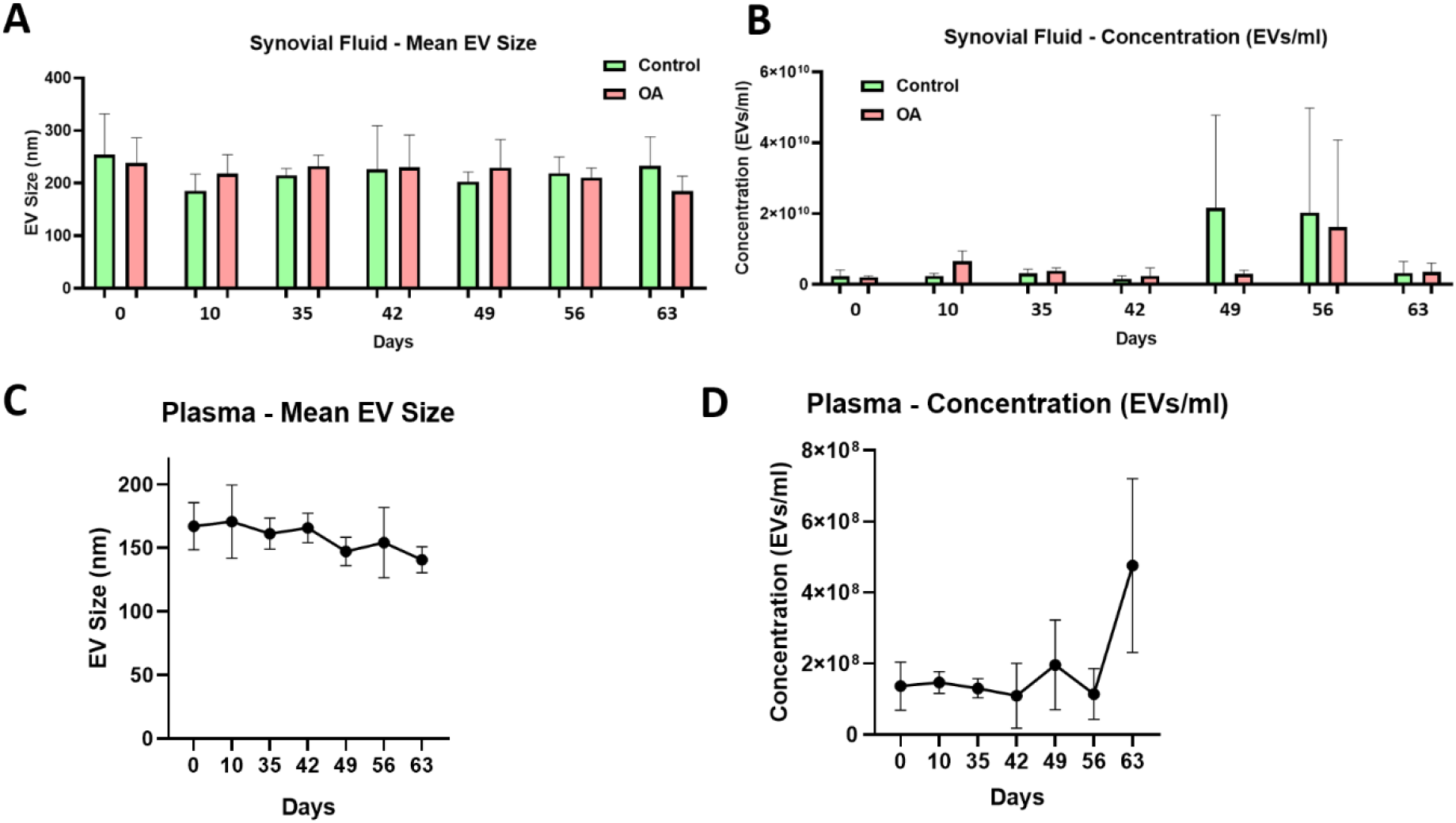
Synovial fluid and plasma derived extracellular vesicle size and concentration. (A) Mean size of extracellular vesicles isolated from synovial fluid control (green) and osteoarthritic (red) middle carpal joints at intervals between 0-63 days following model induction. (B) Mean concentration of isolated extracellular within synovial fluid. (C) Mean size of extracellular vesicles isolated from plasma at intervals between 0-63 days following model induction. (D) Mean concentration of isolated extracellular within plasma. All analyses were conducted via nanoparticle tracking using a Nanosight NS300. Error bars ± 1 standard deviation.

#### Exoview

We analysed the vesicle count, size and heterogeneity of SF and plasma EVs using Exoview in order to validate the particles isolated were EVs. The human ExoView chip assessed CD9, CD63 and CD81markers. For all equine samples there was minimal binding on CD63 capture suggesting low sequence homology between human and equine CD63. CD63 data was therefore excluded from further analysis. Particles stained for CD9 and CD81 were more obvious in plasma-derived EVs (Figure 2A) due to the higher concentration of EVs compared to SF (approximate factor of 10 times lower in SF-derived EVs) (Figure 3A). Plasma EVs had mean diameters between 57-67 nm, (Figure 2B, 2C) and SF 62-74nm (Figure 3B, 3D) both were dependant on the tetraspanin subpopulation. The diameter of the temporal population of plasma-derived EVs changed significantly for both CD9 (Figure 2C) and CD81 (Figure 2D) following OA induction. The number of plasma-derived EVs peaked significantly at day 42 for both CD9 (Figure 2D) and CD81 (Figure 2E) following OA induction. Interestingly in SF there was no difference in vesicle diameter with time or OA for CD9 (Figure 3B). However, there were differences in temporal and disease related expression of CD81 captured vesicles in SF (Figure 3C). There were statistical differences in temporal expression of CD9 vesicles in control but not OA samples (Figure 3D). For CD9 and CD81 in control samples only, expression was lowest at day 42 (Figure 3D, 3E). The expression of CD81 labelled vesicles was significantly altered at day 0 and day 42 in SF (Figure 3E). From the colocalisation results of CD9 and CD81 data there was an apparent drop off in the proportion of EVs double positive at the final time point, day 63 in plasma (Supplementary File 3).

**Figure 2.**
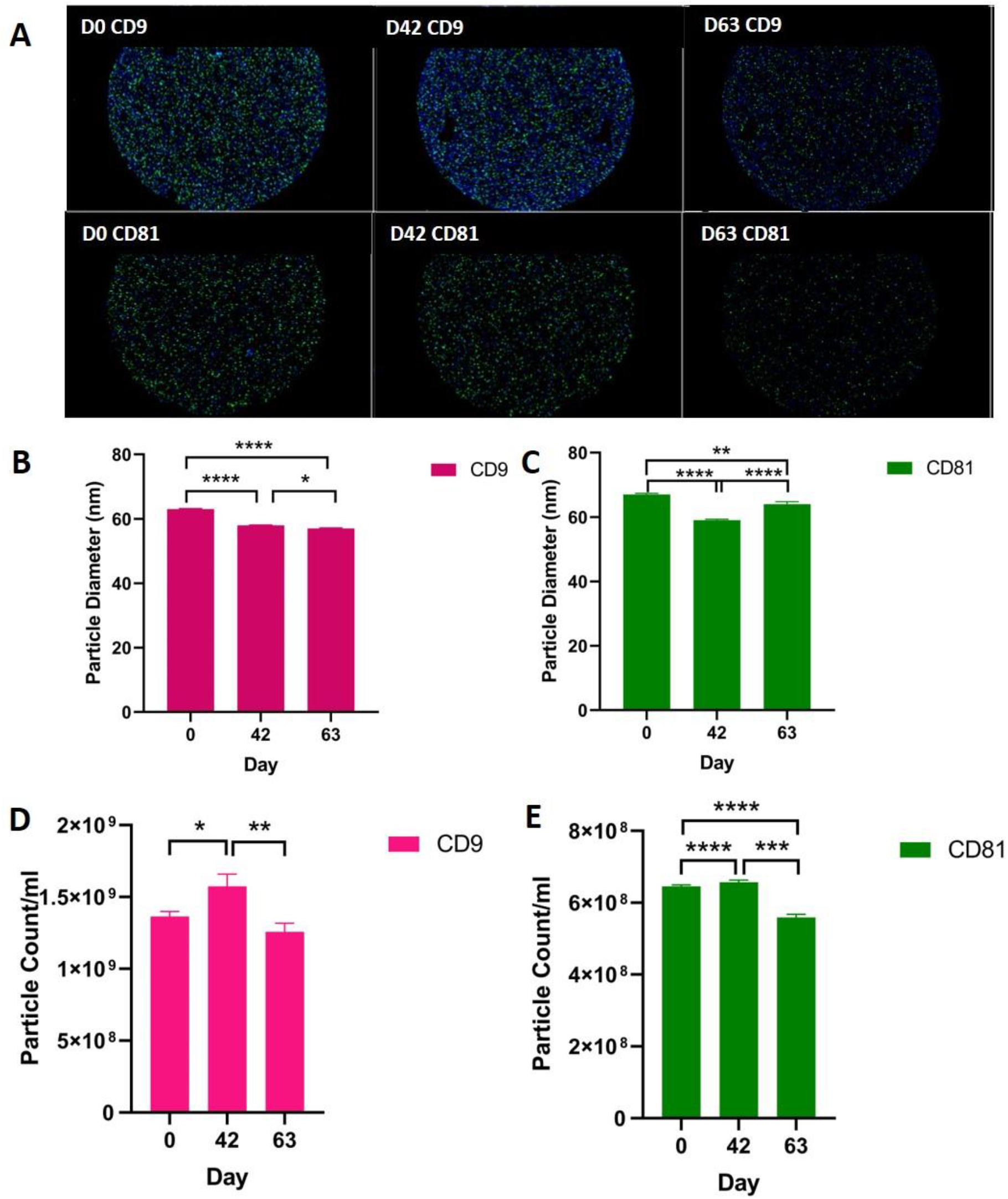
Visualisation, sizing and enumeration of plasma derived EVs. All data was adjusted for dilution of the sample onto the chip. Average of three technical replicates that were run. Particle number was quantified by the number of particles in a defined area on the antibody capture spot. All bars are mean and error bars standard error mean. (A) Fluorescent image of a representative spot for each sample. (B, C) Sizing of CD9 (B) and CD81 (C) labelled EVs, normalized to MIgG control. Limit of detection was 50-200 nm. (D, E) Counting of CD9, and CD81-positive particles after probing with fluorescent tetraspanin antibodies. Statistical analysis undertaken in GraphPad Prism 9.0 using T-tests following parametric evaluation (P<0.05, *; P<0.01 **: p<0.001, ***, p<0.0001, ****).

**Figure 3.**
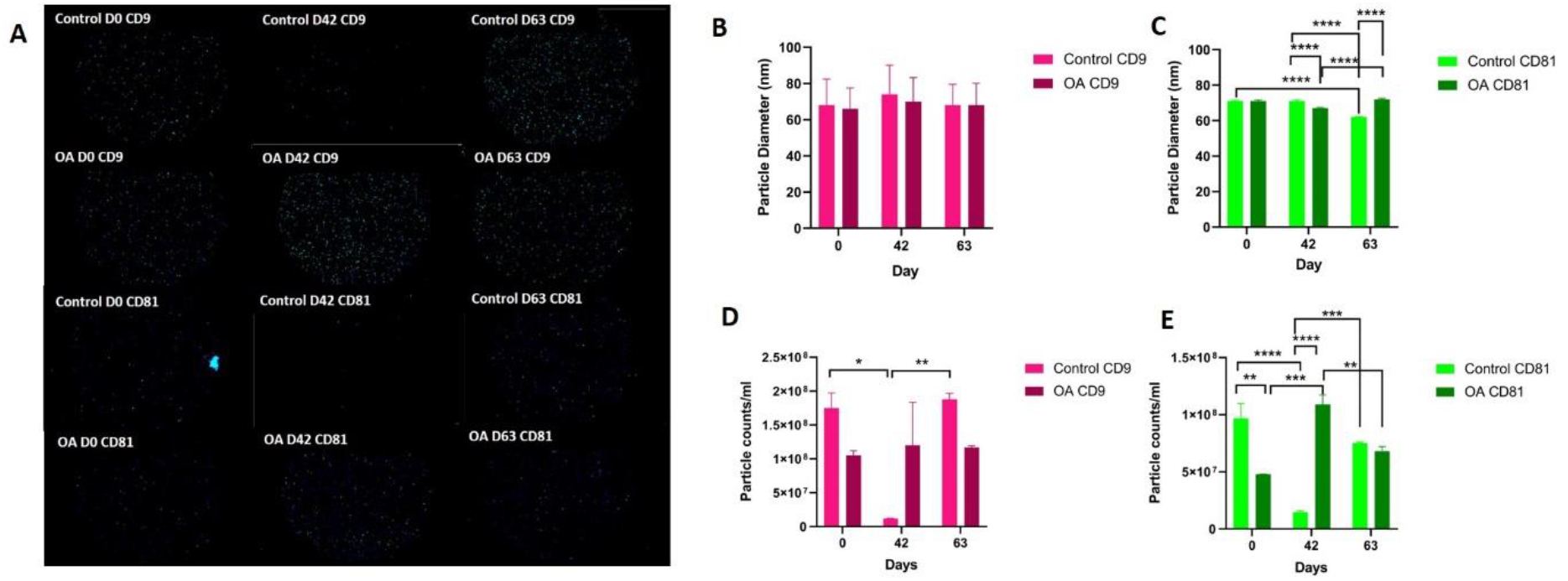
Visualisation, sizing and enumeration of SF derived EVs from control and OA joints. All data was adjusted for dilution of the sample onto the chip. Average of three technical replicates that were run. Particle number was quantified by the number of particles in a defined area on the antibody capture spot. All bars are mean and error bars standard error mean. (A) Fluorescent image of a representative spot for each sample. (B, C) Sizing of CD9 (B) and CD81 (C) labelled EVs, normalized to MIgG control. Limit of detection was 50-200 nm. (D, E) Counting of CD9 (D), and CD81 (E) positive particles after probing with fluorescent tetraspanin antibodies. Statistical analysis undertaken in GraphPad Prism 9.0 using repeated measures anova (P<0.05, *; P<0.01 **: p<0.001, ***, p<0.0001, ****).

### Small RNAseq results

In SF and plasma-derived EVs we identified multiple classes of non-coding RNAs including miRs, snoRNAs, small nuclear RNAs (snRNAs), transfer RNAs (tRNAs), long non-coding RNAs (lncRNA), y-RNA, piwi RNAs (piRNAs) and scaRNAs (scRNA) (Figure 3A). MiRs) were the highest percentage of total reads in plasma whereas in SF it was tRNAs. Supplementary File 4 shows the number of each class of sncRNAs that were identified in at least 30% of samples within that group.

#### MicroRNAs

First, we analysed the changes in plasma miRs before and after OA induction by comparing day 0 to each subsequent time point (days 10, 35, 42, 49, 56, 63) (Table 1) and found 16 different miRs DE. Following pairwise statistical analysis of all-time point plasma-derived EVs, we identified 27 miRs DE (Table 2A, Figure 4B, 4C). Five of these were identified in multiple pairwise comparisons (miR-215, miR-34a, let-7c, miR-130a, miR-146a) (Figure 5). When SF-derived EVs from control or OA joints were interrogated we identified 45 miRs DE (Figure 4D, 4E, Supplementary File 5). Of these 27 were altered between OA timepoints or between control (sham) and OA, representing the most likely miRs involved in OA pathogenesis (Table 2B, Figure 6). Ten of these were identified in multiple pairwise comparisons (miR-let7c, miR-10b, miR-21, miR-25, miR-26a, miR-451, miR-486-5p, miR-744, miR-8993, miR-92a) (Figure 6). A total of seven miRs were DE in plasma and SF (when differences between control and OA or OA at different time points was accounted for); miR-451, miR-25, miR-215, miR-92a, miR-let-7c, miR-486-5p, miR-23a. Figure 7 summarises the changing miR landscape in longitudinal samples for plasma (time only; Figure 7A) and SF (time and disease; Figure 7B).

**Table 1.** Differentially expressed miRNAs compared to day 0 in plasma derived EVs

| Comparison | miRNA | logFC | PValue |
| --- | --- | --- | --- |
| d0 vs d 10 | eca-miR-182 | -11.225 | 0.011 |
| d0 vs d 10 | eca-miR-376c | 4.297 | 0.026 |
| d0 vs d 10 | eca-miR-144 | 5.090 | 0.034 |
| do vs d35 | eca-miR-34a | 11.390 | 0.003 |
| do vs d35 | eca-miR-215 | 9.711 | 0.005 |
| do vs d35 | eca-let-7g | -10.335 | 0.027 |
| do vs d35 | eca-miR-107b | -5.592 | 0.037 |
| do vs d35 | eca-miR-151-5p | -6.071 | 0.047 |
| do vs d42 | eca-miR-24 | -6.513 | 0.034 |
| do vs d49 | eca-miR-23a | 3.996 | 0.011 |
| do vs d49 | eca-miR-1468 | 9.710 | 0.018 |
| do vs d49 | eca-miR-182 | -9.869 | 0.027 |
| do vs d49 | eca-miR-30e | -8.330 | 0.029 |
| do vs d49 | eca-miR-92a | -1.786 | 0.031 |
| do vs d49 | eca-miR-186 | -5.651 | 0.041 |
| do vs d56 | eca-let-7g | -6.541 | 0.031 |
| do vs d56 | eca-miR-19b | -5.617 | 0.031 |
| do vs d56 | eca-miR-151-5p | -6.634 | 0.031 |
| do vs d63 | eca-miR-1307 | 8.836 | 0.014 |

**Table 2A.** Differentially expressed miRNAs following pairwise analysis in plasma derived EVs

| **Pairwise Comparison (Day)** | **miRNA** | **logFC** | **P-Value** | **FDR** |
| --- | --- | --- | --- | --- |
| 35 vs 0 | eca-miR-215 | 7.047 | 0.023 | 0.6559 |
| 35 vs 0 | eca-miR-34a | 8.177 | 0.016 | 0.6559 |
| 35 vs 10 | eca-let-7c | 3.954 | 0.047 | 0.7573 |
| 42 vs 10 | eca-miR-126-5p | 5.043 | 0.033 | 0.4617 |
| 42 vs 10 | eca-miR-141 | -9.149 | 0.005 | 0.2886 |
| 42 vs 10 | eca-miR-144 | -9.702 | 0.017 | 0.4617 |
| 42 vs 10 | eca-miR-215 | -10.11 | 0.030 | 0.4617 |
| 42 vs 35 | eca-let-7c | -2.771 | 0.037 | 0.1648 |
| 42 vs 35 | eca-miR-486-5p | -1.72 | 0.005 | 0.0475 |
| 49 vs 0 | eca-miR-29a | 3.12 | 0.047 | 0.9867 |
| 49 vs 10 | eca-miR-451 | -3.949 | 0.019 | 0.9595 |
| 49 vs 35 | eca-let-7c | -3.903 | 0.005 | 0.04998 |
| 49 vs 35 | eca-miR-92a | -3.685 | 0.004 | 0.04998 |
| 49 vs 42 | eca-miR-23a | 5.417 | 0.031 | 0.9863 |
| 56 vs 35 | eca-miR-215 | -7.294 | 0.034 | 0.5313 |
| 56 vs 35 | eca-miR-25 | -7.204 | 0.045 | 0.5313 |
| 56 vs 35 | eca-miR-34a | -8.436 | 0.028 | 0.5313 |
| 56 vs 35 | eca-miR-99b | -6.102 | 0.036 | 0.5313 |
| 56 vs 42 | eca-miR-130a | 8.764 | 0.000 | 0.02078 |
| 56 vs 42 | eca-miR-23b | 5.075 | 0.017 | 0.4334 |
| 56 vs 49 | eca-miR-130a | 8.579 | 0.006 | 0.3152 |
| 63 vs 35 | eca-miR-107a | 4.513 | 0.049 | 0.5487 |
| 63 vs 35 | eca-miR-181b | 6.285 | 0.047 | 0.5487 |
| 63 vs 35 | eca-miR-215 | -8.491 | 0.013 | 0.5487 |
| 63 vs 35 | eca-miR-34a | -7.571 | 0.033 | 0.5487 |
| 63 vs 42 | eca-miR-146a | 9.981 | 0.045 | 0.9827 |
| 63 vs 56 | eca-miR-146a | 9.609 | 0.041 | 0.818 |

miRNA; microRNA, eca; equine, logFC; log fold change, FDR; false discovery rate

**Table 2B.**
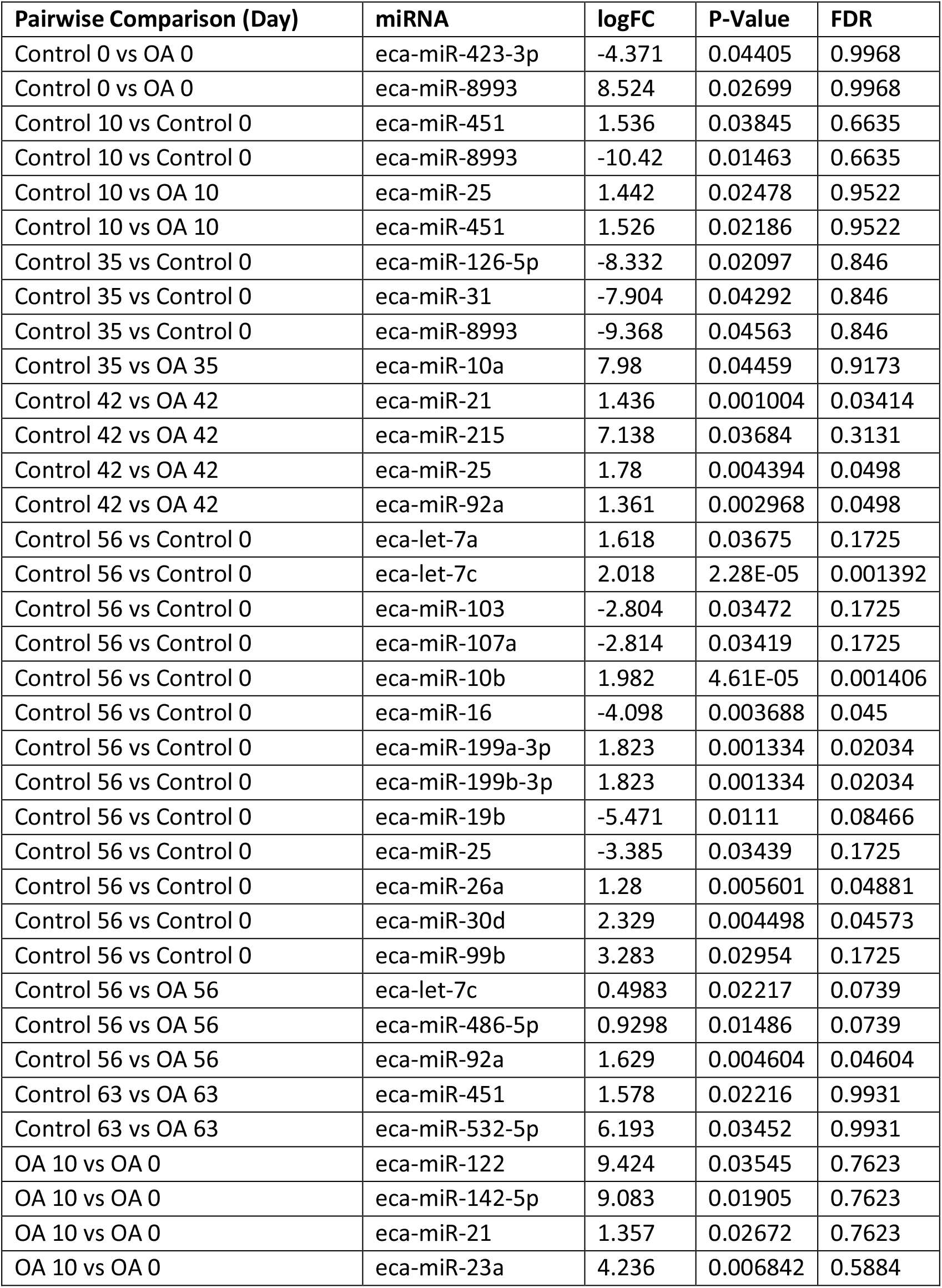
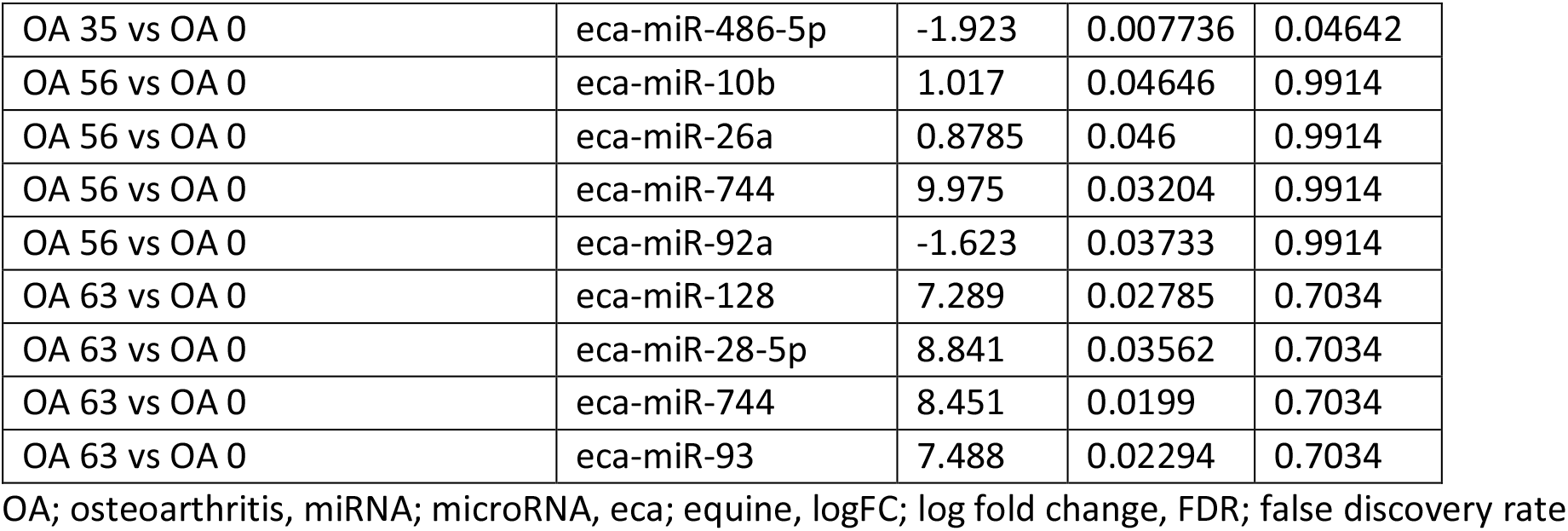
Differentially expressed miRNAs following pairwise analysis in synovial fluid derived EVs

**Table 2B.** Differentially expressed snoRNAs following pairwise analysis in synovial fluid derived EVs

| **Pairwise Comparison (Day)** | **snoRNA name** | **logFC** | **P-Value** | **FDR** |
| --- | --- | --- | --- | --- |
| Control 35 vs Control 0 | U3 | -0.37 | 0.027 | 0.082 |
| OA 35 vs OA 0 | snord27 | -9.9 | 0.038 | 0.094 |
| OA 35 vs OA 0 | snord2 | -9 | 0.043 | 0.094 |
| OA 35 vs OA 0 | snord46 | -9.5 | 0.043 | 0.094 |
| OA 35 vs OA 0 | snord15 | -9.5 | 0.047 | 0.094 |
| OA 49 vs OA 0 | snord27 | -9.8 | 0.023 | 0.095 |
| OA 49 vs OA 0 | snord46 | -9.4 | 0.036 | 0.095 |
| OA 49 vs OA 0 | snord21 | 9.3 | 0.047 | 0.095 |
| OA 56 vs OA 0 | snord46 | -8.9 | 0.043 | 0.093 |
| OA 56 vs OA 0 | snord15 | -8.8 | 0.048 | 0.093 |
| OA 56 vs OA 0 | snord27 | -9.2 | 0.052 | 0.093 |
| OA 56 vs OA 0 | u3 | -1.3 | 0.053 | 0.093 |
| Control 56 v sOA 56 | snord46 | 8.8 | 0.024 | 0.098 |
| Control 63 vs Control 0 | snord58 | 7 | 0.003 | 0.034 |
| OA 10 vs OA 0 | snord58 | 2.3 | 0.019 | 0.097 |
| OA 10 vs OA 0 | snord46 | -3.5 | 0.040 | 0.099 |
| OA 10 vs OA 0 | snord15 | 3 | 0.060 | 0.100 |
| Control 49 vs OA 49 | snord58 | 4.4 | 0.009 | 0.094 |
| Control 56 vs OA 56 | snord15 | -2.9 | 0.000 | 0.000 |
| Control 56 vs OA 56 | snord96 | -2.8 | 0.001 | 0.010 |
| Control 63 vs OA 63 | snord46 | -3.7 | 0.003 | 0.043 |

snRNA; small nucleolar RNA, logFC; log fold change, FDR; false discovery rate

**Figure 4.**
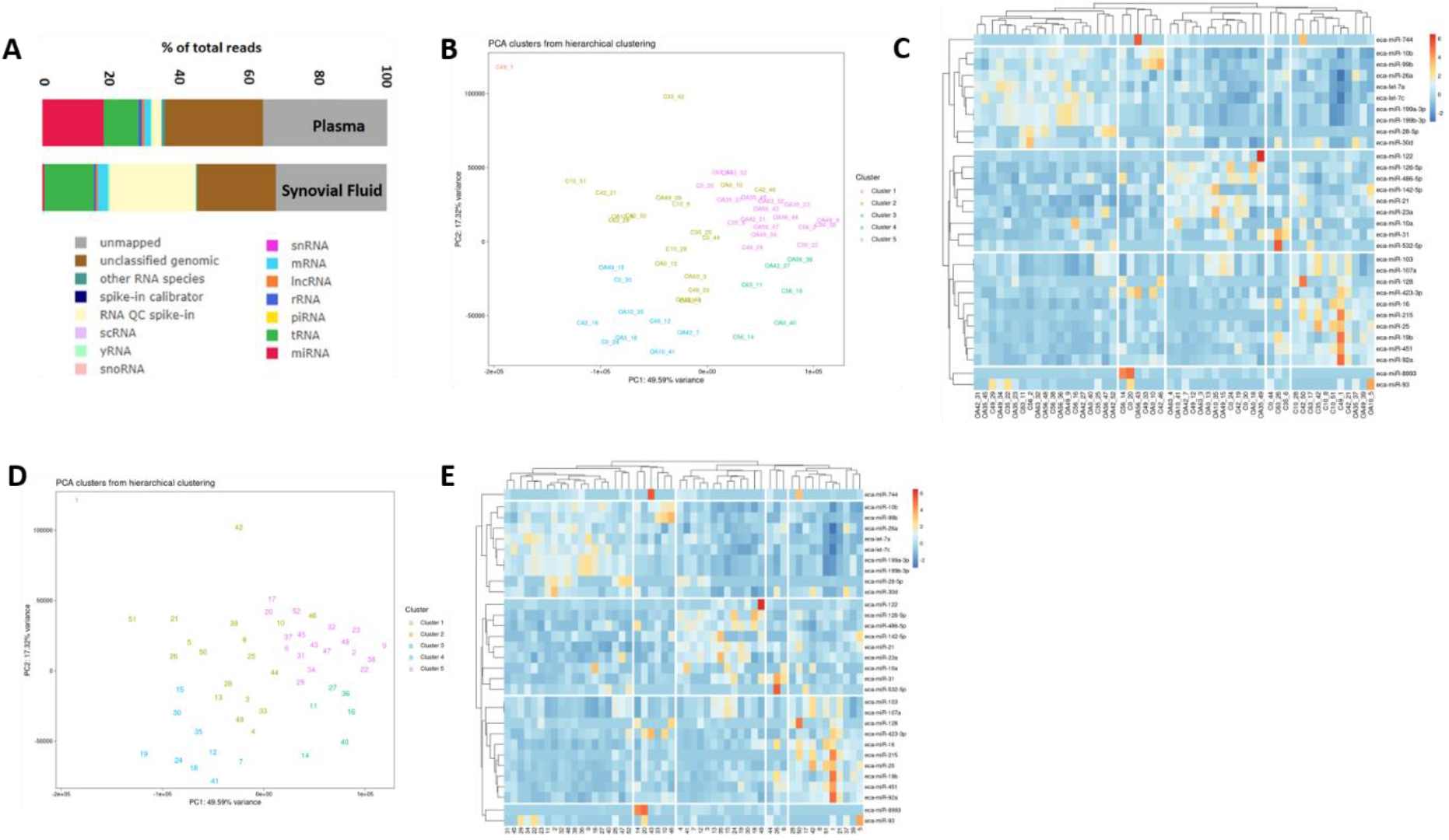
Summary data for small non-coding RNA sequencing of plasma and synovial fluid derived EVs. (A) Total reads in plasma and synovial fluid based on RNA type. (B) PCA of DE microRNAs for plasma derived EVs. (C) Heatmap of differentially expressed microRNAs (P<0.05) in plasma derived EVs. (D) PCA of microRNAs for SF derived EVs; C, control; OA, osteoarthritis. Numbering of samples; the first number relates to day of sampling; 0, 10, 35, 42, 49, 56, 63. The second number is the original sample ID. (E) Heatmap of DE microRNAs (P<0.05) in SF derived EVs. Heatmaps created using the R pheatmap package. Clustering was undertaken with the “average” method and correlation distancing. Scaling is unit variance scaled expression, based on read counts. Clustering distances are based on Pearson correlation.

**Figure 5.**
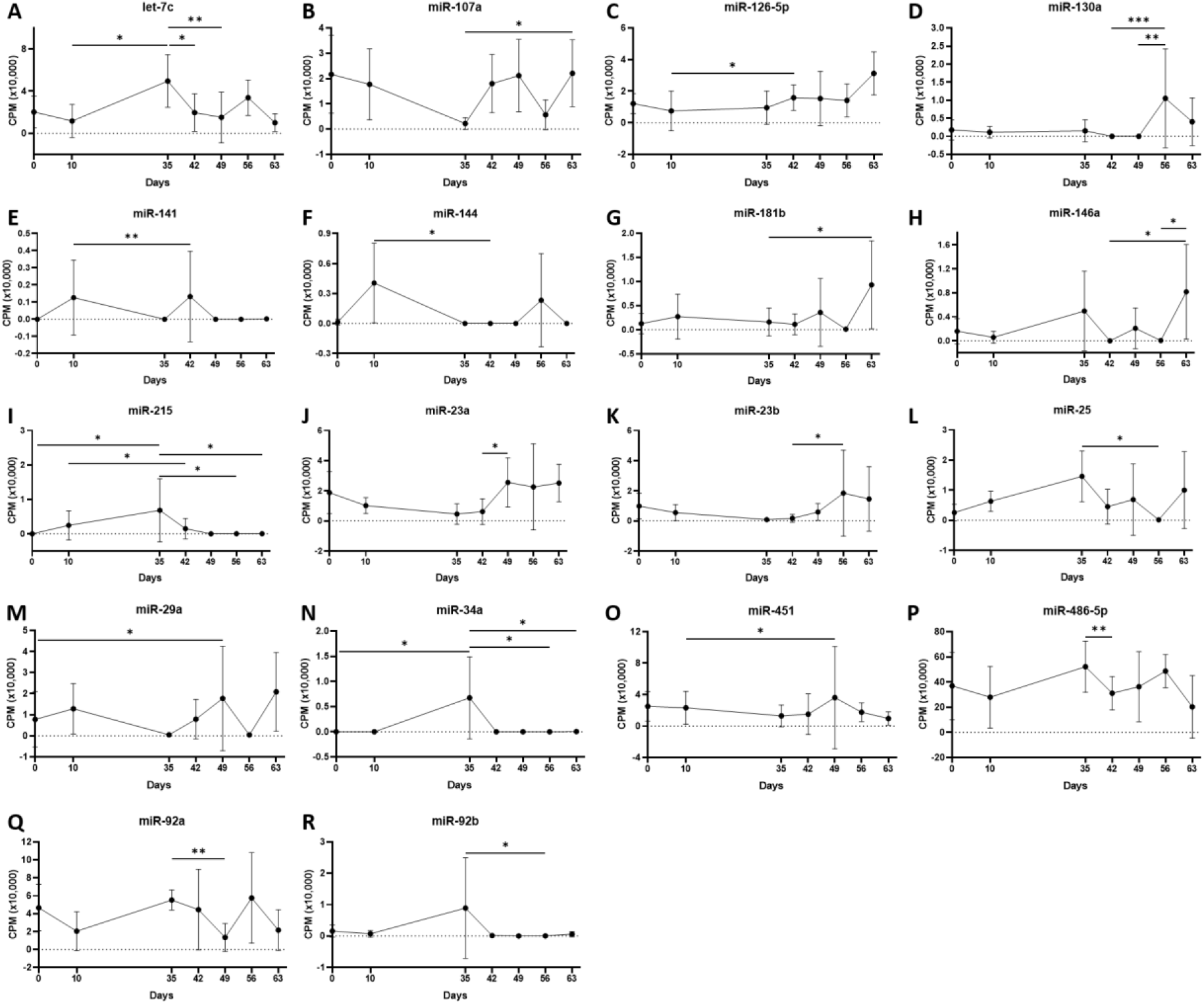
Differentially expressed miRNAs isolated from plasma-derived extracellular vesicles. Individual panels for miRs DE using pairwise comparisons. Error bars ± 1 standard deviation. * = p < 0.05, ** = p < 0.01, and *** = p < 0.001.

**Figure 6.**
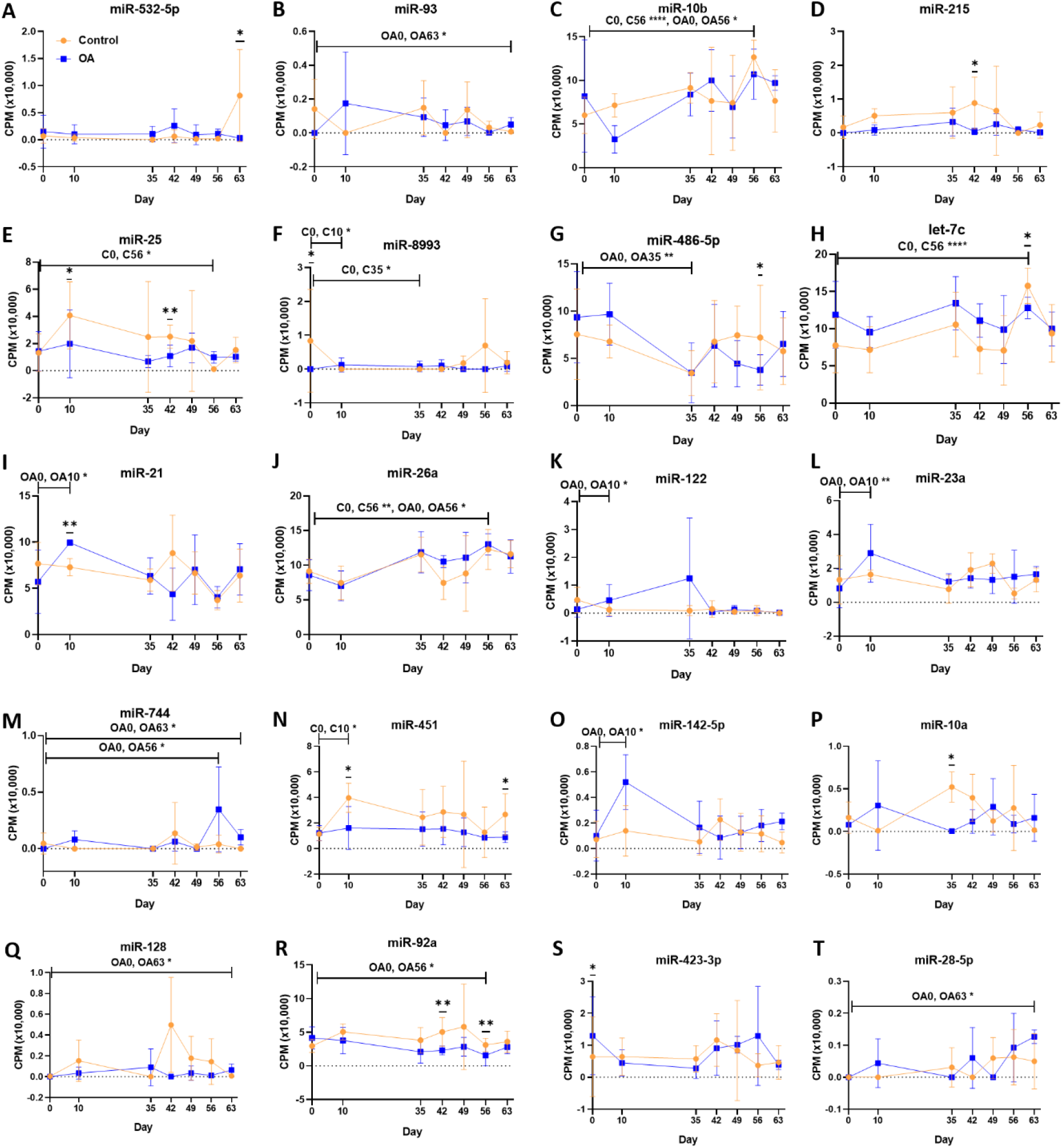
Differentially expressed miRNAs isolated from synovial fluid-derived extracellular vesicles. Comparisons made between control (C) and osteoarthritis (OA) and between OA time points (0,10,35,42,49,56,63). Error bars ± 1 standard deviation. * = p < 0.05, ** = p < 0.01 and **** = p< 0.0001.

**Figure 7.**
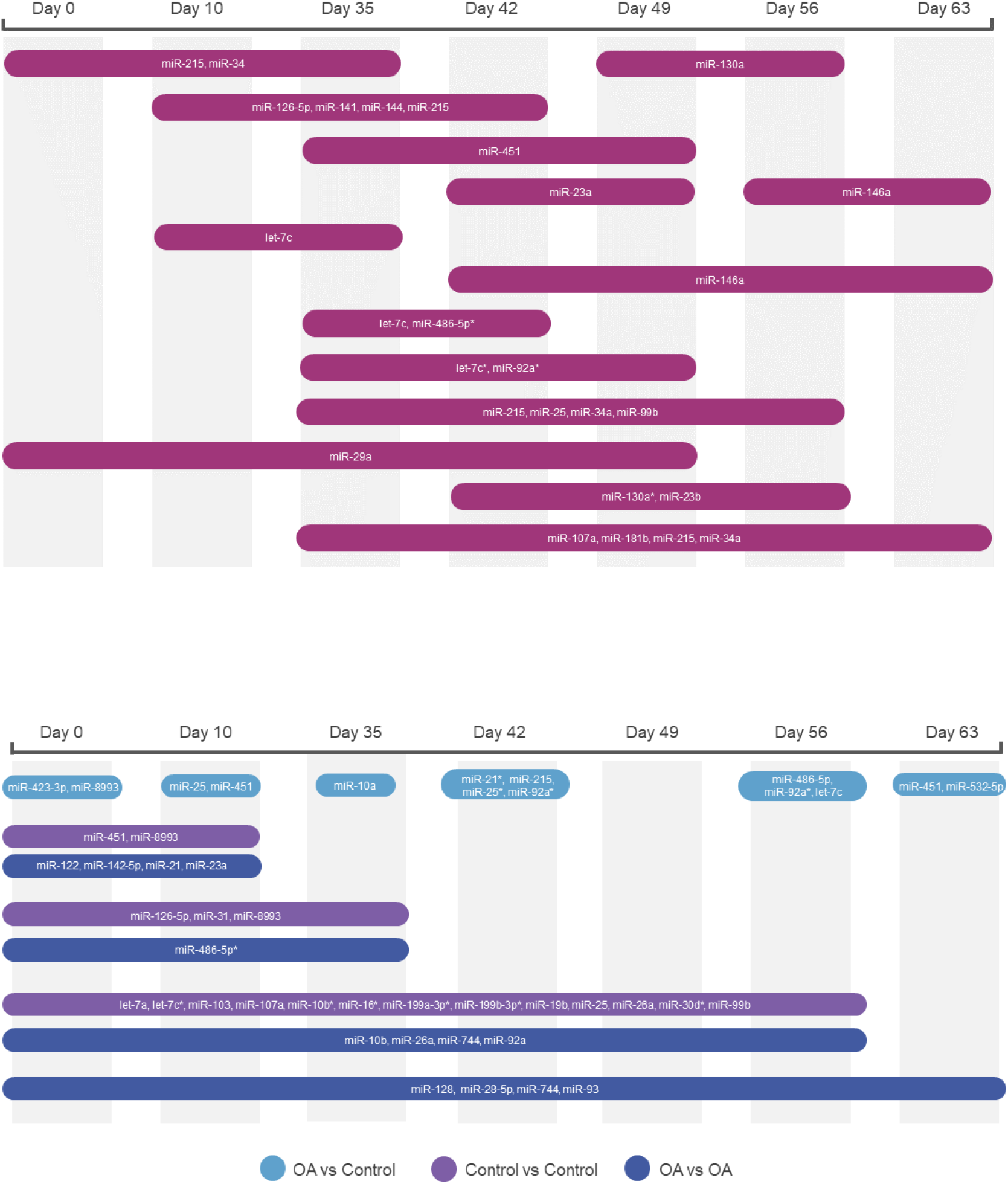
Diagrams summarises the changing miR landscape in longitudinal samples. (A) Plasma timeline (B) Synovial fluid with time and diseases. MiRs identified following pairwise comparisons P<0.05, * denotes FDR<0.05.

#### SnoRNAs

Following pairwise statistical analysis considering donor effect of plasma-derived EVs, we identified 35 snoRNAs DE (Table 3A, Figure 8A, 8B). Ten of these were identified in multiple pairwise comparisons (snord102, U3, snord113, snord113/114, snord15, snord66, snord69, snord58, snord62, snord65). When SF-derived EVs from control or OA joints were interrogated we identified 21 snoRNAs DE (Table 3B, Figure 8C, 8D). Six of these were identified in multiple pairwise comparisons (U3, snord27, snord15, snord46, snord27, snord58). There were four snoRNAs DE in plasma and SF (U3, snord15, snord46, snord58).

**Figure 8.**
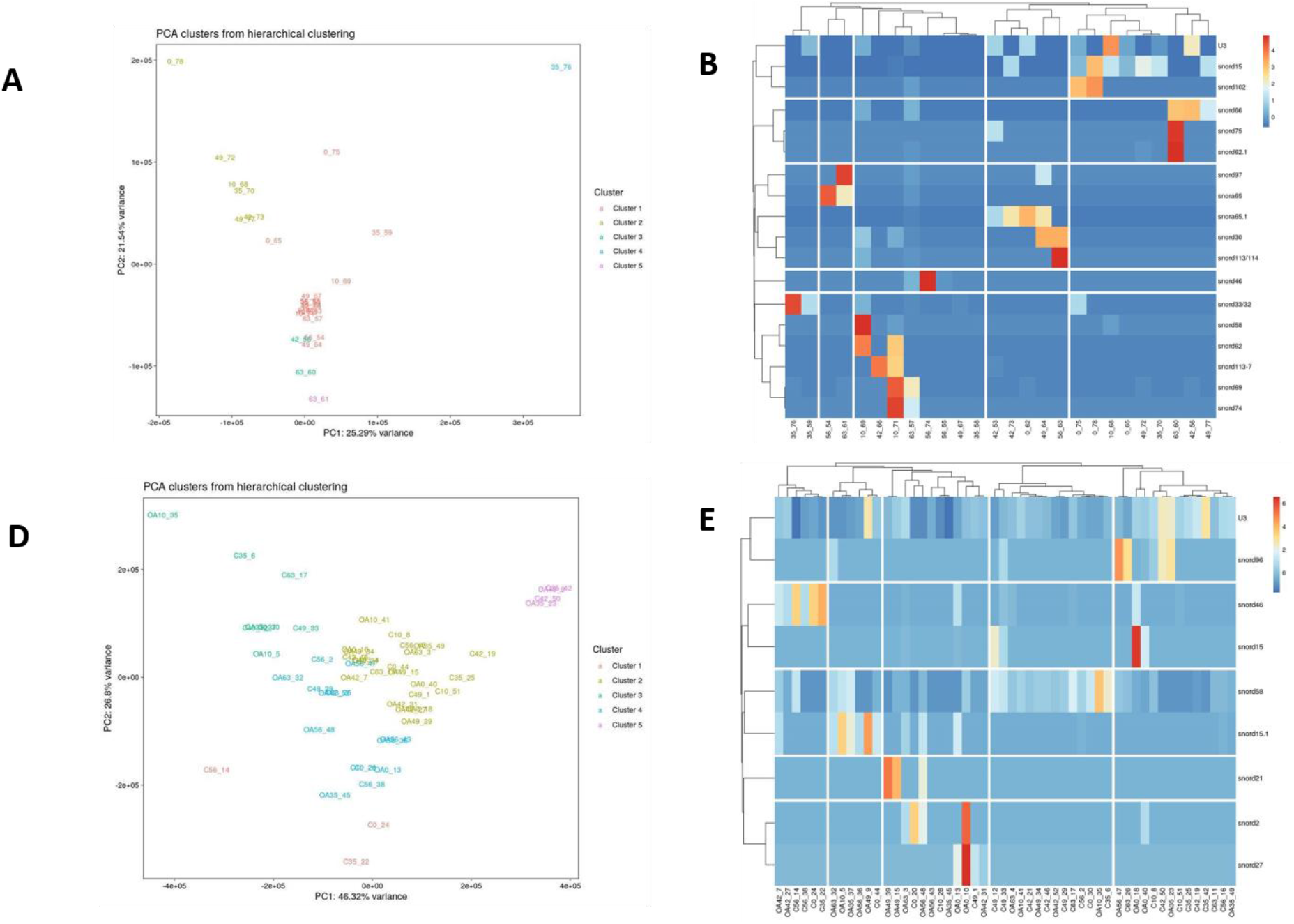
Differentially expressed snoRNAs in plasma and synovial fluid derived EVs. (A) PCA of DE snoRNAs for plasma derived EVs. (C) Heatmap of DE snoRNAs (P<0.05) in plasma derived EVs. (D) PCA of snoRNAs for SF derived EVs; C, control; OA, osteoarthritis. Numbering of samples; the first number relates to day of sampling; 0, 10, 35, 42, 49, 56, 63. The second number is the original sample ID. (E) Heatmap of DE snoRNAs (P<0.05) in SF derived EVs. Heatmaps created using the R pheatmap package. Clustering was undertaken with the “average” method and correlation distancing. Scaling is unit variance scaled expression, based on read counts. Clustering distances are based on Pearson correlation.

**Table 3A.** Differentially expressed snoRNAs following pairwise analysis in plasma derived EVs

| **Pairwise Comparison (Day)** | **snoRNA name** | **logFC** | **P-Value** | **FDR** |
| --- | --- | --- | --- | --- |
| 56 vs 0 | U3 | -6.3 | 0.000 | 0.000 |
| 56 vs 0 | snord102 | -7.8 | 0.000 | 0.002 |
| 56 vs 0 | snord46 | 4.6 | 0.001 | 0.003 |
| 49 vs 42 | snord113-7 | -4.2 | 0.003 | 0.041 |
| 56 vs 49 | snord15 | -6.7 | 0.032 | 0.097 |
| 63 vs 56 | snord66 | 7.4 | 0.009 | 0.092 |
| 49 vs 0 | U3 | -7 | 0.006 | 0.082 |
| 56 vs 0 | snord15 | -7.9 | 0.032 | 0.096 |
| 56 vs 0 | snord102 | -8.2 | 0.048 | 0.096 |
| 63 vs 0 | snord66 | 9.8 | 0.024 | 0.094 |
| 35 vs 10 | snord69 | -9.9 | 0.003 | 0.046 |
| 42 vs 10 | snord113/114 | -3.9 | 0.005 | 0.076 |
| 42 vs 10 | snord69 | -8.2 | 0.013 | 0.076 |
| 42 vs 10 | snord58 | -6.9 | 0.015 | 0.076 |
| 42 vs 10 | snord62 | -7.1 | 0.016 | 0.076 |
| 42 vs 10 | snord30 | -8 | 0.025 | 0.095 |
| 56 vs 10 | snord69 | -11 | 0.002 | 0.026 |
| 56 vs 10 | snord15 | -7.1 | 0.013 | 0.092 |
| 56 vs 10 | U3 | -6.8 | 0.027 | 0.092 |
| 56 vs 10 | snord62 | -7.1 | 0.031 | 0.092 |
| 56 vs 10 | snord58 | -7 | 0.034 | 0.092 |
| 56 vs 10 | snord74 | -8.7 | 0.039 | 0.092 |
| 56 vs 35 | snord33/32 | -9.3 | 0.006 | 0.063 |
| 56 vs 35 | U3 | -8.8 | 0.019 | 0.094 |
| 63 vs 35 | snord66 | 11 | 0.012 | 0.096 |
| 63 vs 35 | snord97 | 11 | 0.012 | 0.096 |
| 63 vs 35 | snord75 | 10 | 0.019 | 0.096 |
| 63 vs 35 | snord113/114 | 9.3 | 0.039 | 0.096 |
| 63 vs 35 | snord69 | 9.2 | 0.039 | 0.096 |
| 63 vs 35 | snord62 | 9 | 0.039 | 0.096 |
| 63 vs 35 | snora65 | 8 | 0.039 | 0.096 |
| 56 vs 42 | U3 | -7.7 | 0.011 | 0.080 |
| 56 vs 42 | snora65 | -8.4 | 0.028 | 0.099 |
| 56 vs 49 | snord15 | -7.8 | 0.029 | 0.086 |

snRNA; small nucleolar RNA, logFC; log fold change, FDR; false discovery rate

#### Other non-coding RNAs

Pairwise comparisons for other non-coding RNAs considering donor effect were undertaken in plasma and SF. In plasma we found five tRNAs, eight lncRNAs, and five snRNAs DE (FDR<0.1) Additionally, in SF we identified five tRNAs, two lncRNAs, and four snRNAs DE (FDR<0.1) (Supplementary File 6).

### Bioinformatics analysis of DE miRs in plasma or SF

#### Plasma pairwise comparison

For plasma-derived EVs first we looked at the set of 16 DE miRs following pairwise comparisons compared to day 0. IPA identified the top diseases and functions with negative activated Z scores (used to infer the activation states of biological functions based on comparison with a model that assigns random regulation directions; for negative this relates to a prediction of inhibition) as migration (P=9.14 E-04), cell proliferation (P=5.32 E-05) and cell viability (P=8.7 E-02) (Figure 9A). To investigate the position of the 27 DE miRNA plasma expression network, we then determined their putative target genes using IPA Target Filter which integrates computational algorithms with multiple miRNA databases. We used a threshold of ‘experimentally validated’ or ‘highly predicted’. This matched the miRs to 920 mRNA targets. The mRNA targets were input into the gene ontology tool ToppGene, and then, the 1626 biological processes identified were visualised using Revigo and Cytoscape (Figure 9B)). The top 100 biological processes were visualised in a Treemap (Figure 9B).

**Figure 9.**
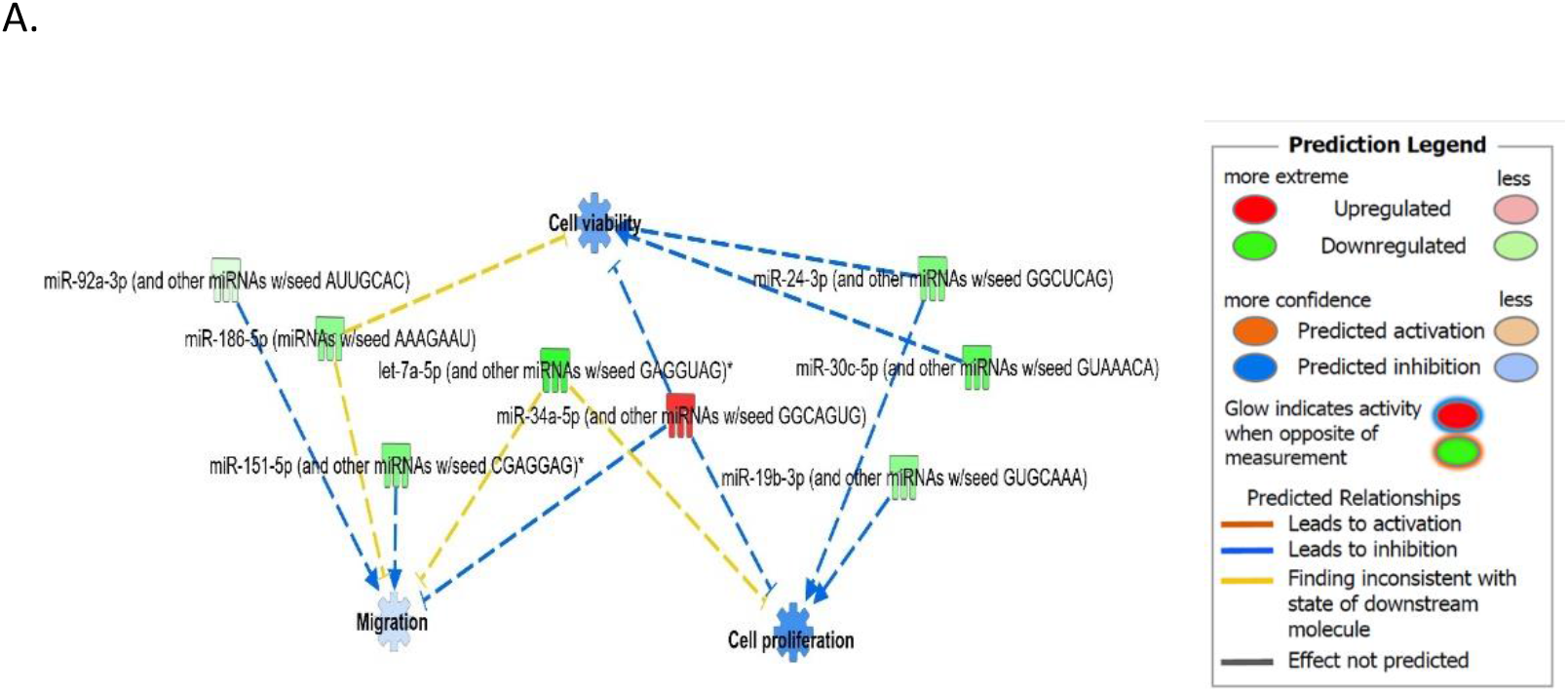
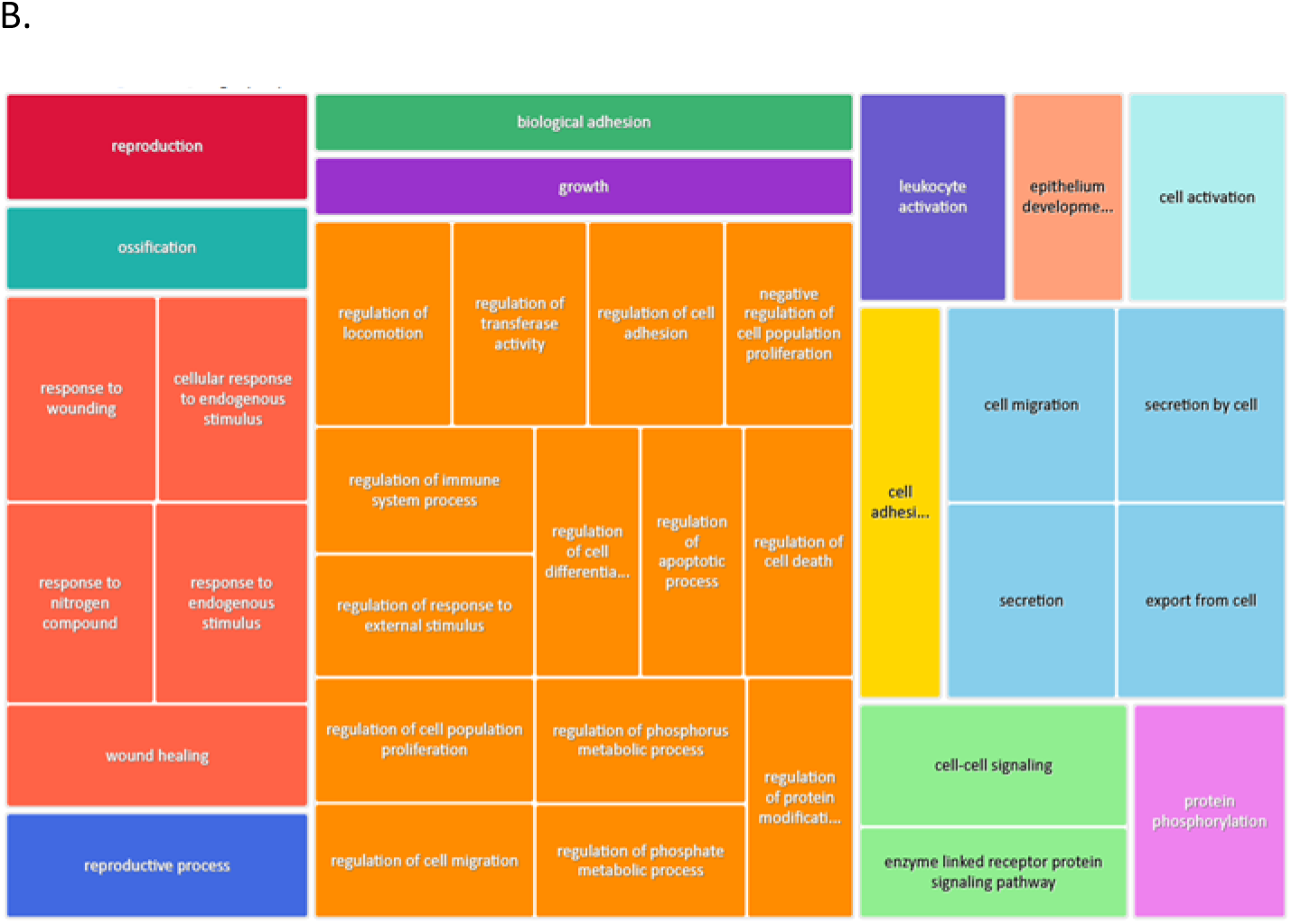
Bioinformatics analysis of the DE plasma derived EV miR and their putative mRNA targets. A. IPA of the 27 DE Mirs following all pairwise comparisons compared to day 0 showed cell viability, proliferation and migrations were activated. B. A treemap of the top 100 GO terms. GO biological processes associated with dysregulated miRNA targets were identified following TargetScan filter module in IPA. ToppGene was used to perform functional enrichment analysis on predicted miRNA targets to highlight biological processes most significantly affected by dysregulated miRNA-mRNA interactions. GO terms (FDR < 0.05) were summarized and visualised using REViGO and Cytoscape. Allowed similarity setting in Revigo was medium.

#### SF pairwise comparisons

We input for bioinformatics analysis only miRs DE in SF pairwise comparisons altered between control (sham) and OA or between OA samples at different timepoints. Thus 27 miRs were input into IPA ‘Core Analysis’ (Supplementary File 7). The transcription factors E2F Transcription Factor 1 (P=7.37 E-07)and 3 (3.68E-09) and Argonaute RISC Catalytic Component 2 (p=3.75 E-17), an initiator of target mRNAs degradation were significant upstream regulators of eight DE miRs with downstream effects on senescence (p=3.66E-04), inflammation (p=2.32E-03), apoptosis (p=7.57 E-04), angiogenesis (p=1.17E-05), fibrosis (6.84E-16), gene silencing (p=1.74E-10), transcription (p=0.002) and expression of RNA (p=0.003) (Figure 10A). A heatmap derived from IPA identified biological processes based on activation Z-score (Figure 10B). This highlighted that cell cycle was predicted to be inhibited (negative activation Z-score) and contributed to by G1 phase (p=1.55E-08, Z-score −1.6), G1/S phase transition (p=3.52E-06, Z-score −1.6), interphase (p=6.45E-08, Z-score-1.5), senescence (p=3.7E-04, Z-score −0.67) and cell cycle progression (p=1.5 E-05, Z-score −0.66) (Figure 10C, Supplementary File 6). Furthermore, whilst DNA damage (p=1.32E-04, Z-score-2) and cell proliferation (p=9.06E-07, Z-score −2.2) were also reduced, cell viability (p=1.6E-03, Z-score 0.75), differentiation of stem cells (p=2.36E-04, Z-score 1.03) were predicted to be increased (Figure 10C).

**Figure 10.**
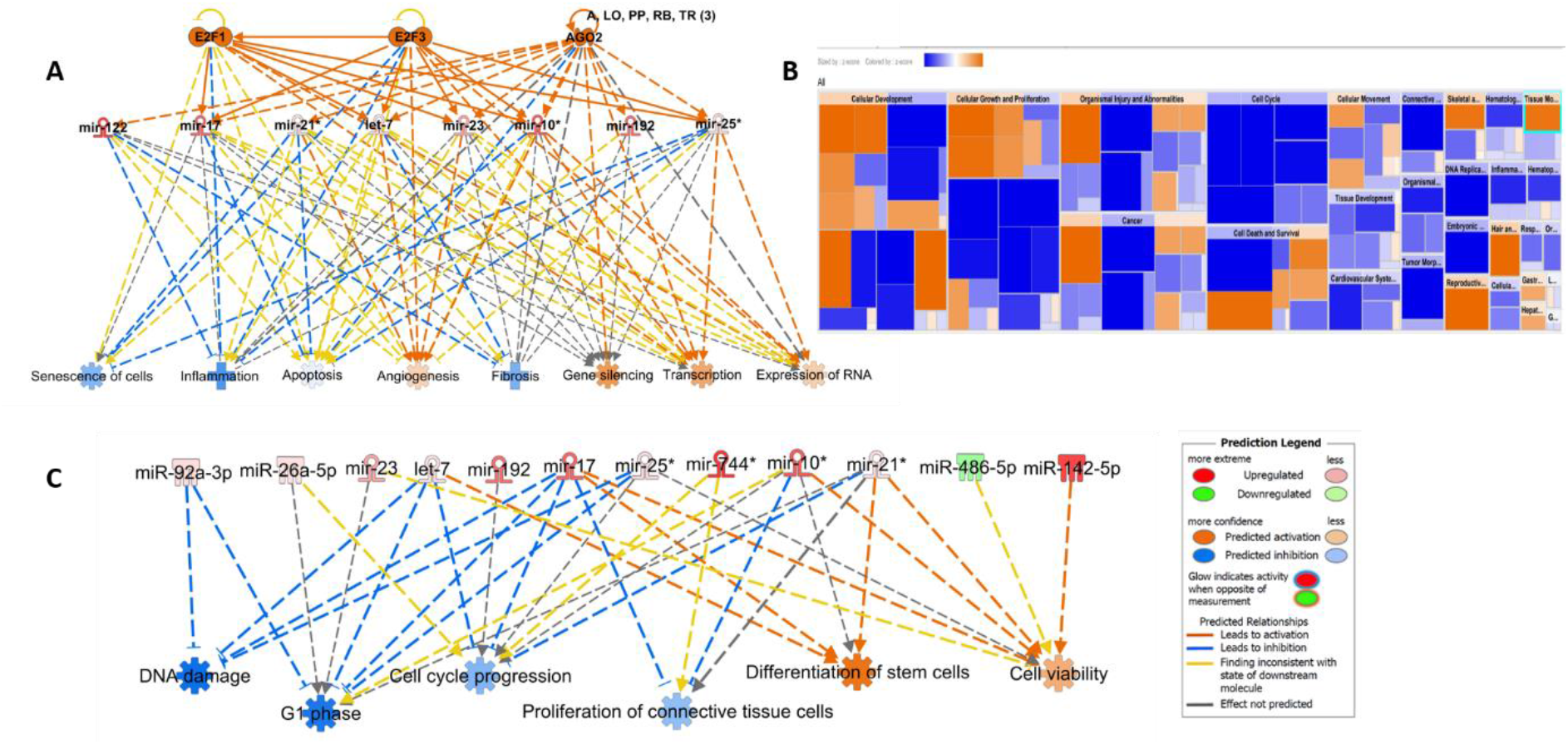
Pathway analysis of miRs DE in synovial fluid using IPA. (A) Regulatory effects network for E2F1, E2F3 and AGO2 actions senescence, inflammation, apoptosis, angiogenesis, fibrosis, gene silencing, transcription and expression of RNA via the differentially expressed miRNAs. (B) Heatmap sized by z score of biological processes with scale were blue is inhibited and orange activated. (C) Significant diseases and functions based on Z-score associated with DE miRs. Figures are graphical representations of molecules identified in our data in their respective networks. Red nodes; upregulated and green nodes; downregulated gene expression. Intensity of colour is related to higher fold-change. Legends to the main features in the networks are shown. The biological function is dependent on whether it is predicted to be activated or inhibited.

## Discussion

Fundamental studies that contribute to characterising and understanding early OA will serve to elucidate disease pathogenesis, identify novel molecules as biomarkers for early detection or act as therapeutic targets. The demonstration that EVs are a method of cell communication has caused a recent increase in EV publications (49, 50). In this study we have, for the first time, studied the sncRNA EV content temporally in OA using an equine model. In our previous work we have confirmed the development of OA in our equine carpal osteochondral fragment model using histological evaluation. There was significant cartilage degradation and synovial membrane inflammation in OA joints (48).

Similarities in the pathogenesis and clinical signs of OA in man and horse are leading the expansion of equine translational studies (59, 60). As well as the advantages of histological and anatomical similarities to human joints it is easy to undertake longitudinal sampling of SF and blood. However, the use of the horse as a model is limited by high costs and ethical considerations which reduces the number of animals in studies. In our experiment we had access to samples from four horses but were able to examine, for the first-time in any species, temporal OA changes in EV characteristics and sncRNA cargo using SF and plasma and spanning seven time points from day 0-72 (study end). Whilst our groups have examined the sncRNA cargo in equine SF in early OA (8) and the temporal changes in microRNAs in SF from same equine osteochondral fragment model (39) using less time points here we tried to identify the changing sncRNA landscape within SF or plasma-derived EVs.

It is challenging for most experimental designs to discriminate exosomes from microvesicles in terms of their size, cargo, properties and origin (51). Consequently, most research represents the characteristics of a heterogenous group of nanovesicles which are derived from both endosomal multivesicular bodies and plasma membrane budding. In our study we used both NTA, as well as interferometric images (Exoview) using tetraspanin antibodies to size and count individual EVs. First, we characterised the EV profile in all the samples of SF and plasma from the model. To isolate our EVs we used size exclusion chromatography to separate EVs from proteins and lipids as we had only small volumes of our biological fluids. Others have recommended this isolation methods when there are limited volumes of plasma and SF, and downstream analysis includes sncRNA and protein cargo (52, 53). Ultracentrifugation alone in SF-derived EVs increased contamination and aggregation whilst size exclusion chromatography increased EV enrichment (54).

EVs are involved in multiple functional roles in joint homeostasis (reviewed (55)). EV characteristics of size and concentration can be affected by any external stimuli which significantly alters formation, release and uptake of EVs (56). Using NTA we measured overall sizes and quantity of isolatable EVs. This method enables acceptable repeatability and reproducibility (57). However, the results must be interpreted considering inherent biological sample diversity. The samples contained a heterogenous group of EVs of various sizes and are comparable only with caution to preparations from cell culture and other plasma or SF studies as results are also dependant on methods of EV isolation. Having utilised size exclusion chromatography our measurements displayed minimal levels of background noise reflecting the reduction of protein aggregates or cellular components compared to ultracentrifugation isolation methods (53). Our NTA size estimates were within EV range and similar to other plasma (58) and SF (54, 59) studies using size exclusion chromatography. However compared to a recent plasma EV study in human OA using EV precipitation and nanoparticle tracking the equine plasma EVs were smaller (human 235nm, equine 158nm) (23) which may be due to isolation methods used. Comparing both biofluids the measured particle sizes were overall greater in SF compared to plasma cases. We did not find any difference in EV concentration, size, mode, D10, D50 or D90 in either SF or plasma temporally or with OA induction in the model. Interestingly there are few studies characterising size and concentration of EVs in SF or plasma in normal and OA samples. Our results concur with one study in which NTA was used to characterise EVs in human normal and OA SF (59). However, it should be noted that a precipitation exosome kit was used in this study for EV isolation.

Following exosome isolation from human knee SF there was increase in exosome concentration from early to severe knee OA without changes in the average particle size (60). In a further study of early and late stage knee OA following isolation of exosomes with ultracentrifugation of biofluids, whilst for plasma there was no difference in concentration of exosomes, in SF the expression of exosomes in early OA and late-stage OA was higher than that in controls (61).

As NTA does not specifically measure EVs but also co-isolated contaminants we also used Exoview™ to characterise SF and plasma-derived EVs at selected time points. This technology uses single particle interferometric reflectance imaging sensing together with antibody-based chip capture and fluorescence detection that enables single vesicle identification. Additionally, it has a lower size limit of detection compared to scattering based techniques (62). We had previously tested EVs isolated from equine SF and plasma on both the mouse and human tetraspanin chips and found that the human chips captured EVs more efficiently (data not shown). Whilst classical EV markers are CD9, CD63 and CD81 some cells do not express CD63 (63) and so a panel of antibodies is preferable. There was minimal staining for CD63 indicating either poor antibody sequence homology or rareness of these EVs. As we suspected a lack of sequence homology we did not analyse CD63 results further. Due to prohibitive costs we only ran a subset of the SF and plasma samples on the Exoview platform. In plasma there was an apparent change in size following OA induction. In SF no temporal or OA-related change was evident for CD9, however for CD81 there was. CD81 expression changed with time in SF. The differences in tetraspanin expression may indicate that the EVs present in SF and plasma could have different functional activities as tetraspanins are important for the functionalities of EVs as their function is dependent on their ability to interact with target cells. These in turn are determined by surface receptors existing in each EV subtype (64). Tetraspanin complexes are the EV surface receptors that would define the target cells to bind (65). A study in seminal plasma demonstrated an association of CD63 with CD9 and CD81 in exosomes, indicating a possible synergistic effect of these tetraspanins (66). Simultaneous immunostaining enabled dual immunofluorescent analysis and therefore the co-localisation of CD9 and CD81 was assessed. In plasma at day 0 and day 42 there were similar colocalisation profiles, but the proportion of double positive EVs reduced significantly at day 63. This could potentially indicate a switch in tetraspanin phenotype in OA. The biological implications of this switch are unknown. One limitation of our study which should be noted with respect to Exoview data is the limited number of samples analysed and further work is necessary before hard conclusions can be drawn.

Since the identification of RNA within EVs (67) there has been a rise in the interest of using EV-RNA as biomarkers (68). Others have demonstrated that the relative concentrations of miRs secreted in EVs differs from those found in the cell, or serum/plasma concentrations of miRs (69). Due to the role of EVs in cell communication we hypothesized that sncRNA changes due to OA may be found in both SF but also plasma EVs. SF-derived EVs are likely to come from a number of sources including cartilage, subchondral bone, synovium and the circulation. We used a non-biased approach; small RNA sequencing to remove bias and whilst most blood (20, 22, 70–72) or SF studies (20, 21) in OA have looked at miRs alone we investigated a number of non-coding RNAs in our samples. To our knowledge this is only the second study to investigate EVs derived from plasma in OA (23) and the first to describe SF-derived EV sncRNAs in OA. In plasma we found 16 different miRs DE between day 0 and other time points and 27 DE when all pairwise comparisons were undertaken. We would presume changes in the plasma could be due to the OA model but also be attributed to the sham procedure in which the opposite joint was subject to arthroscopy. Of the miRs DE some have been found to change in OA in other studies; miR-126, miR-29 (22), and miR-146a in four human studies (20, 22, 71, 73). Interestingly whilst one study in plasma found no miRs DE in plasma-derived EVs in OA, miR-146a was one of the most highly expressed miRs identified (23). In our study we demonstrated that it significantly increased between day 42-63 and day 56-63. Mir-146a has a vital role in maintaining cartilage homeostasis and OA (74, 75). In a mouse study surgically induced OA treated with a miR-146a inhibitor significantly alleviated cartilage destruction via targeting Camk2d and Ppp3r2 (74). Serum miR-146a-5p was significantly increased in women with knee OA compared with controls (73).

In SF-derived EVs we identified 27 miRs that were DE between control and OA or temporally in OA. Some have been previously identified as DE in other studies including let7c, miR-10a, miR-122, miR-215 and miR-23a in SF derived from horses 28 days after the osteochondral model commenced in a larger cohort of horses (48).

Seven miRs were DE in plasma and SF-derived EVs; miR-451, miR-25, miR-215, miR-92a, miR-let-7c, miR-486-5p, miR-23a. Thus, these miRs represent the most promising biomarkers for further work. An ideal biomarker for early equine OA would be sourced from blood rather than SF, but the fact that changes occur in both biofluids suggest they are associated with OA changes within our model. Interestingly miR-215, let7-c and miR-23a were also identified in our study in which we isolated sncRNAs from SF. They were altered at day 28 compared to day 0 in the same model and the same horses (48). MiR-23a was also increased in equine early OA SF (8) and increased in late versus early human OA SF (21). Mir-23a contributes to OA progression by directly targeting SMAD3 (76). Generally our miR findings support our other work with this model in which we found that OA disease progression caused early changes in SF miR expression patterns at day 28 (77).

We can only compare the plasma-derived EV results with one previous study at a single non-temporal study in OA in which no DE microRNAs were identified. The study used 23 OA patients and 23 controls and isolated EVs using precipitation followed by sequencing the samples on a similar sequencing platform to that used in this study (23). These discrepancies may be related to differences in species, sample type, disease stage or severity, methodology, gender differences, and other factors among studies.

In order to understand the potential role of the panel of miRs changing in our model we conducted pathway analysis of the DE miRs. Firstly, we took DE plasma-derived miRs following pairwise comparisons compared to day 0. This was to define changes at a pathway level temporally compared to prior to OA induction (day 0). A ‘Core Analysis’ of the miRs in IPA described a predicted inhibition of migration, cell proliferation and cell viability. The carpal osteochondral fragment model specifically simulates a post-traumatic OA phenotype. Therefore, every event from carpal chipping until erosion of cartilage is considered part of the disease pathogenesis. This suggests that plasma-derived miR EV cargo changing in the model would inhibit cell migration, proliferation and viability. However, the origin of these EVs cannot be confirmed but if related to the model would represent changes occurring in both the OA and sham joint including processes including wound healing and repair, bone, cartilage and synovial damage, and reparative processes of these tissues plus inflammation. In OA mesenchymal stem cells migrate and proliferate into the damaged cartilage area (78) and inflammatory cells can migrate to the joint (79). By reducing migration, proliferation and viability the plasma miR EV cargo could potentially affect the interplay of the various cells in the OA joint.

Next, we took the 27 DE miRNA plasma expression network, used IPA to predict their target genes and then condensed the list of biological pathways affected by these mRNAs. This identified many terms related to cell response (including would healing, and endogenous stimuli), regulation of cellular processes (including cell death, proliferation, apoptosis, migration and metabolic processes), plus cell signalling, secretion and activation. As the EV-miR cargo were from plasma it is difficult to distinguish if these effects are due to OA changes alone or also opening both the OA and sham joints. Despite this our findings potentially offer in insight into the early changes in post-traumatic OA. However, by interrogating the potential pathways of the DE miRs in SF (which we hypothesise would be more likely to be due to disease induction) we detailed the potential role of this cargo and its effects in early post traumatic OA in the joint. Interestingly whilst one would expect Argonaute RISC Catalytic Component to be an upstream regulator of the mIRs (as the Argonaute family of proteins have a role in RNA interference (80)), transcription factors E2F Transcription Factor 1 and 3 were also predicted. This has a role in accelerating cell proliferation and promoting inflammation signalling (81) and E2F1 targets may be affected in OA pathogenesis (82).

Additionally, we analysed the expression changes of other sncRNAs. To our knowledge for the first time in an OA model we have identified changing plasma and SF-derived EV tRNA, lncRNAs and snRNAs. These could be important molecules for understanding and treating OA. For example, Liu *et al.* showed that exosomes derived from human mesenchymal stem cells contained lncRNA KLF3-AS1 and promoted chondrocyte proliferation *in vitro.* Whilst in an *in vivo* collagenase-induced OA model these exosomes improved cartilage repair (83).

We also investigated the changes in snoRNAs. From our previous work we know snoRNAs have an important role in joint homeostasis (84), in mouse joint ageing (16), in chondrogenic differentiation (85), in cartilage ageing (14), in OA (6, 7) and in early OA SF (8). Interestingly in plasma we found that 82% of the snoRNAs DE in plasma were also identified in ageing/OA human cartilage study (7), including Snord113/114, snord15, snord30, snord32/33, snord46, snord58, snord62, snord66, snord69, snord74, snord75 and U3. We have also shown snord113 to be DE in ageing mouse joints (16), ageing equine chondrocytes (86), ageing equine cartilage (87); U3 to be DE in ageing mouse serum (16), OA cartilage and chondrocytes (6); snord32/33 in ageing mouse serum (16) and OA human and equine cartilage where it has a role in oxidative stress (88). For DE snoRNAs in synovial fluid 88% of the snoRNAs DE in plasma were also identified in ageing/OA human cartilage study (7) including snord15, snord2, snord21, snord46, snord58, snord96A, U3. Furthermore, snord96A was DE in equine SF in early OA (8) and snord58 in ageing human cartilage (unpublished work). Four snoRNAs; U3, snord15, snord46, snord58 were found DE in both plasma and SF-derived EVs and were all altered in pairwise comparisons in OA. Changes in U3 are particularly interesting given our knowledge of its role in OA and impact on protein translational apparatus (6). This snoRNA was reduced in plasma at multiple time points and in SF between day 0 and day 56. It is essential for rRNA maturation, acting as a spliceosome during ribosome biogenesis, releasing 18S rRNA from the precursor 47S rRNA (89). Impaired 18S rRNA is suggested to further decrease levels of 5.8S and 28S rRNA in experimental U3 impaired human articular chondrocytes (6). Whilst in OA human articular chondrocytes following U3 downregulation, there was a reduction in expression of chondrogenic genes, accompanied by increased levels of chondrocyte hypertrophy genes. OA-related U3 downregulation effects the protein translation capacity of chondrocytes *in vitro* (6). Additionally, sequences of U3 have been suggested to have miRNA-like abilities, accomplishing effective RNA-silencing, *in vitro,* in at least two human cell lines (89). Overall the changes in snoRNAs in EVs in both plasma and SF provide insight into their role in the pathogenesis of OA as traditionally OA has been defined as an imbalance between joint anabolism/catabolism. With our increasing evidence that it is an acquired ribosomopathy (84), there is a potential role for EV-derived snoRNAs to contribute to OA pathogenesis via its cell communication role. We are currently studying sncRNA cross talk in joint cells to shed more light on this hypothesis (90). Thus, EV snoRNAs may provide novel molecular opportunities for the development of OA therapeutics.

We realise there are a number of limitations in the study. The plasma and SF samples used in this study were stored at −80°C for two years prior to analysis and we were limited in the volumes of plasma and SF we had due to multiple platforms samples were used for. Although sample collection, handling and storage was the same for all samples any inconsistencies in these procedures may alter the levels of sncRNAs. We used spike-ins throughout the sequencing methodology to increase reproducibility, and indeed the number of sncRNAs was similar to those observed by others in plasma-derived EVs (23) suggesting our results are likely to be representative for sncRNAs in SF and plasma EVs. Unfortunately, we were unable to validate our findings in the same samples using qRT-PCR as there was not enough remaining sample for EV extraction.

## Conclusion

Sequencing of temporal samples for sncRNAs in an equine model produced unbiased profiling of the circulating and SF-derived EV sncRNAome and identified a unique panel of sncRNAs in during initiation and progression of early post-traumatic equine OA. We characterised plasma and SF-derived EVs in equine OA for the first time and demonstrated that differences in tetraspanin expression may indicate that in early OA they could represent changing functionalities of EVs. The DE of seven miRs DE in plasma and SF-derived EVs; miR-451, miR-25, miR-215, miR-92a, miR-let-7c, miR-486-5p, miR-23a and four snoRNAs; U3, snord15, snord46, snord58 represent exciting molecules for future work.

## Acknowledgments

This study and James Anderson were funded by the Horserace Betting Levy Board. Mandy Peffers was funded through a Wellcome Trust Clinical Intermediate Fellowship (grant 107471/Z/15/Z). This work was also supported by the Independent Research Fund Denmark, Technology and Production Sciences [grant number DFF - 7017-00066, 2017], the Horse Levy foundation, and Gerda and Aage Haensch’s Foundation. Marie Walters was funded by a PhD scholarship jointly awarded by the University of Copenhagen, the Technical University of Denmark and the Swedish University of Agricultural Science. The authors thank Catarina Castanheira for help with figure production and Marianne Pultar for help with additional data analysis.

## Author contributions

Conception and design of the animal experiment: MW, LB, STJ

Collection of data: JRA, MJP, EJC, MW, LB, STJ

Analysis and interpretation of data: JRA, MJP, EJC, AD, MH, VJ, MW, STJ

Statistical analysis and expertise: JRA, MJP, AD, MH

Drafting the article: MJP, JRA

Obtaining of funding; STJ, MW, MJP.

Critical revision of the article for important intellectual content: All authors

Final approval of the article: All authors

## References

1. Hunter DJ, Bierma-Zeinstra S. Osteoarthritis. Lancet. 2019;393(10182):1745–59.

2. Ireland JL, Clegg PD, McGowan CM, McKane SA, Chandler KJ, Pinchbeck GL. Comparison of owner-reported health problems with veterinary assessment of geriatric horses in the United Kingdom. Equine Vet J. 2012;44(1):94–100.

3. Todhunter PG, Kincaid SA, Todhunter RJ, Kammermann JR, Johnstone B, Baird AN, et al. Immunohistochemical analysis of an equine model of synovitis-induced arthritis. Am J Vet Res. 1996;57(7):1080–93.

4. Little CB, Ghosh P, Rose R. The effect of strenuous versus moderate exercise on the metabolism of proteoglycans in articular cartilage from different weight-bearing regions of the equine third carpal bone. Osteoarthritis Cartilage. 1997;5(3):161–72.

5. Goldring MB. Update on the biology of the chondrocyte and new approaches to treating cartilage diseases. Best Pract Res Clin Rheumatol. 2006;20(5):1003–25.

6. Ripmeester EGJ, Caron MMJ, van den Akker GGH, Surtel DAM, Cremers A, Balaskas P, et al. Impaired chondrocyte U3 snoRNA expression in osteoarthritis impacts the chondrocyte protein translation apparatus. Sci Rep. 2020;10(1):13426.

7. Peffers MJ, Chabronova A, Balaskas P, Fang Y, Dyer P, Cremers A, et al. SnoRNA signatures in cartilage ageing and osteoarthritis. Sci Rep. 2020;10(1):10641.

8. Castanheira C, Balaskas P, Falls C, Ashraf-Kharaz Y, Clegg P, Burke K, et al. Equine synovial fluid small non-coding RNA signatures in early osteoarthritis. BMC Vet Res. 2021;17(1):26.

9. Choudhuri S. Small noncoding RNAs: biogenesis, function, and emerging significance in toxicology. J Biochem Mol Toxicol. 2010;24(3):195–216.

10. Tonge DP, Pearson MJ, Jones SW. The hallmarks of osteoarthritis and the potential to develop personalised disease-modifying pharmacological therapeutics. Osteoarthritis Cartilage. 2014;22(5):609–21.

11. Fu H, Hu D, Zhang L, Tang P. Role of extracellular vesicles in rheumatoid arthritis. Mol Immunol. 2018;93:125–32.

12. Sluijter JPG, Davidson SM, Boulanger CM, Buzas EI, de Kleijn DPV, Engel FB, et al. Extracellular vesicles in diagnostics and therapy of the ischaemic heart: Position Paper from the Working Group on Cellular Biology of the Heart of the European Society of Cardiology. Cardiovasc Res. 2018;114(1):19–34.

13. Besse B, Charrier M, Lapierre V, Dansin E, Lantz O, Planchard D, et al. Dendritic cell-derived exosomes as maintenance immunotherapy after first line chemotherapy in NSCLC. Oncoimmunology. 2016;5(4):e1071008.

14. Peffers MJ, Liu X, Clegg PD. Transcriptomic profiling of cartilage ageing. Genom Data. 2014;2:27–8.

15. Balaskas P, Goljanek-Whysall K, Clegg P, Fang Y, Cremers A, Emans P, et al. MicroRNA Profiling in Cartilage Ageing. Int J Genomics. 2017;2017:2713725.

16. Steinbusch MM, Fang Y, Milner PI, Clegg PD, Young DA, Welting TJ, et al. Serum snoRNAs as biomarkers for joint ageing and post traumatic osteoarthritis. Sci Rep. 2017;7:43558.

17. Castanheira C, Anderson JR, Fang Y, Milner PI, Goljanek-Whysall K, House L, et al. Mouse microRNA signatures in joint ageing and post-traumatic osteoarthritis. Osteoarthr Cartil Open. 2021;3(4):100186.

18. Miyaki S. [Cartilage/chondrocyte research and osteoarthritis. The role of microRNAs and extracellular vesicles in osteoarthritis pathogenesis.]. Clin Calcium. 2018;28(6):783–8.

19. Miyaki S, Lotz MK. Extracellular vesicles in cartilage homeostasis and osteoarthritis. Curr Opin Rheumatol. 2018;30(1):129–35.

20. Murata K, Yoshitomi H, Tanida S, Ishikawa M, Nishitani K, Ito H, et al. Plasma and synovial fluid microRNAs as potential biomarkers of rheumatoid arthritis and osteoarthritis. Arthritis Res Ther. 2010;12(3):R86.

21. Li YH, Tavallaee G, Tokar T, Nakamura A, Sundararajan K, Weston A, et al. Identification of synovial fluid microRNA signature in knee osteoarthritis: differentiating early-and late-stage knee osteoarthritis. Osteoarthritis Cartilage. 2016;24(9):1577–86.

22. Borgonio Cuadra VM, Gonzalez-Huerta NC, Romero-Cordoba S, Hidalgo-Miranda A, Miranda-Duarte A. Altered expression of circulating microRNA in plasma of patients with primary osteoarthritis and in silico analysis of their pathways. PLoS One. 2014;9(6):e97690.

23. Aae TF, Karlsen TA, Haugen IK, Risberg MA, Lian OB, Brinchmann JE. Evaluating plasma extracellular vesicle microRNAs as possible biomarkers for osteoarthritis. Osteoarthritis and Cartilage Open. 2020.

24. Kaur S, Abu-Shahba AG, Paananen RO, Hongisto H, Hiidenmaa H, Skottman H, et al. Small non-coding RNA landscape of extracellular vesicles from human stem cells. Sci Rep. 2018;8(1):15503.

25. Ni Z, Zhou S, Li S, Kuang L, Chen H, Luo X, et al. Exosomes: roles and therapeutic potential in osteoarthritis. Bone Res. 2020;8:25.

26. Withrow J, Murphy C, Liu Y, Hunter M, Fulzele S, Hamrick MW. Extracellular vesicles in the pathogenesis of rheumatoid arthritis and osteoarthritis. Arthritis Res Ther. 2016;18(1):286.

27. Ragni E, Perucca Orfei C, De Luca P, Lugano G, Vigano M, Colombini A, et al. Interaction with hyaluronan matrix and miRNA cargo as contributors for in vitro potential of mesenchymal stem cell-derived extracellular vesicles in a model of human osteoarthritic synoviocytes. Stem Cell Res Ther. 2019;10(1):109.

28. Cai J, Wu J, Wang J, Li Y, Hu X, Luo S, et al. Extracellular vesicles derived from different sources of mesenchymal stem cells: therapeutic effects and translational potential. Cell Biosci. 2020;10:69.

29. Teeple E, Jay GD, Elsaid KA, Fleming BC. Animal models of osteoarthritis: challenges of model selection and analysis. AAPS J. 2013;15(2):438–46.

30. Frisbie DD, Kawcak CE, Trotter GW, Powers BE, Walton RM, McIlwraith CW. Effects of triamcinolone acetonide on an in vivo equine osteochondral fragment exercise model. Equine Vet J. 1997;29(5):349–59.

31. McIlwraith CW, Frisbie DD, Kawcak CE. The horse as a model of naturally occurring osteoarthritis. Bone Joint Res. 2012;1(11):297–309.

32. Frisbie DD, Kawcak CE, McIlwraith CW. Evaluation of the effect of extracorporeal shock wave treatment on experimentally induced osteoarthritis in middle carpal joints of horses. Am J Vet Res. 2009;70(4):449–54.

33. McIlwraith CW, Frisbie DD, Kawcak CE, Fuller CJ, Hurtig M, Cruz A. The OARSI histopathology initiative - recommendations for histological assessments of osteoarthritis in the horse. Osteoarthritis Cartilage. 2010;18 Suppl 3:S93–105.

34. Filipe V, Hawe A, Jiskoot W. Critical evaluation of Nanoparticle Tracking Analysis (NTA) by NanoSight for the measurement of nanoparticles and protein aggregates. Pharm Res. 2010;27(5):796–810.

35. Andrews. S. FastQC: A quality control tool for high throughput sequence data https://www.bioinformatics.babraham.ac.uk/projects/fastqc/2010 [

36. Ewels P, Magnusson M, Lundin S, Kaller M. MultiQC: summarize analysis results for multiple tools and samples in a single report. Bioinformatics. 2016;32(19):3047–8.

37. Martin. M. Cutadapt removes adapter sequences from high-throughput sequencing reads. EMBnetjournal. 2011;17(1):10.

38. Langmead B, Trapnell C, Pop M, Salzberg SL. Ultrafast and memory-efficient alignment of short DNA sequences to the human genome. Genome Biol. 2009;10(3):R25.

39. Friedlander MR, Mackowiak SD, Li N, Chen W, Rajewsky N. miRDeep2 accurately identifies known and hundreds of novel microRNA genes in seven animal clades. Nucleic Acids Res. 2012;40(1):37–52.

40. Zerbino DR, Achuthan P, Akanni W, Amode MR, Barrell D, Bhai J, et al. Ensembl 2018. Nucleic Acids Res. 2018;46(D1):D754–D61.

41. S G-J. The microRNA Registry. Nucleic Acids Research. 2004;32:109D–11.

42. Blake Sweeny A, Petrov AI, Burkov B, Finn R, D.,, Bateman A, Szymanski M, et al. RNAcentral: A hub of information for non-coding RNA sequences. Nucleic Acids Research 2019;47:D221–29.

43. Robinson MD, McCarthy DJ, Smyth GK. edgeR: a Bioconductor package for differential expression analysis of digital gene expression data. Bioinformatics. 2010;26(1):139–40.

44. Love MI, Huber W, Anders S. Moderated estimation of fold change and dispersion for RNA-seq data with DESeq2. Genome Biology 2014;15(12):1–21.

45. Chen J, Bardes EE, Aronow BJ, Jegga AG. ToppGene Suite for gene list enrichment analysis and candidate gene prioritization. Nucleic Acids Res. 2009;37(Web Server issue):W305–11.

46. Supek F, Bosnjak M, Skunca N, Smuc T. REVIGO summarizes and visualizes long lists of gene ontology terms. PLoS One. 2011;6(7):e21800.

47. Shannon P, Markiel A, Ozier O, Baliga NS, Wang JT, Ramage D, et al. Cytoscape: a software environment for integrated models of biomolecular interaction networks. Genome Res. 2003;13(11):2498–504.

48. Walters M, Skovgaard K, Heegaard P, Peffers MJ, Fang Y, Bundgaard L, et al. Changes in small non-coding RNA expression in synovial fluid during disease progression in an equine model of experimental osteoarthritis. Osteoarthritis and Cartilage. 2021;29:SS155.

49. Harding CV, Heuser JE, Stahl PD. Exosomes: looking back three decades and into the future. J Cell Biol. 2013;200(4):367–71.

50. Ale Ebrahim S, Ashtari A, Zamani Pedram M, Ale Ebrahim N, Sanati-Nezhad A. Publication Trends in Exosomes Nanoparticles for Cancer Detection. Int J Nanomedicine. 2020;15:4453–70.

51. Raposo G, Stahl PD. Extracellular vesicles: a new communication paradigm? Nat Rev Mol Cell Biol. 2019;20(9):509–10.

52. Tzaridis T, Bachurski D, Liu S, Surmann K, Babatz F, Gesell Salazar M, et al. Extracellular Vesicle Separation Techniques Impact Results from Human Blood Samples: Considerations for Diagnostic Applications. Int J Mol Sci. 2021;22(17).

53. Brennan K, Martin K, FitzGerald SP, O’Sullivan J, Wu Y, Blanco A, et al. A comparison of methods for the isolation and separation of extracellular vesicles from protein and lipid particles in human serum. Sci Rep. 2020;10(1):1039.

54. Foers AD, Chatfield S, Dagley LF, Scicluna BJ, Webb AI, Cheng L, et al. Enrichment of extracellular vesicles from human synovial fluid using size exclusion chromatography. J Extracell Vesicles. 2018;7(1):1490145.

55. Esa A, Connolly KD, Williams R, Archer CW. Extracellular Vesicles in the Synovial Joint: Is there a Role in the Pathophysiology of Osteoarthritis? Malays Orthop J. 2019;13(1):1–7.

56. Zaborowski MP, Balaj L, Breakefield XO, Lai CP. Extracellular Vesicles: Composition, Biological Relevance, and Methods of Study. Bioscience. 2015;65(8):783–97.

57. Vestad B, Llorente A, Neurauter A, Phuyal S, Kierulf B, Kierulf P, et al. Size and concentration analyses of extracellular vesicles by nanoparticle tracking analysis: a variation study. J Extracell Vesicles. 2017;6(1):1344087.

58. Holcar M, Ferdin J, Sitar S, Tusek-Znidaric M, Dolzan V, Plemenitas A, et al. Enrichment of plasma extracellular vesicles for reliable quantification of their size and concentration for biomarker discovery. Sci Rep. 2020;10(1):21346.

59. Kolhe R, Hunter M, Liu S, Jadeja RN, Pundkar C, Mondal AK, et al. Gender-specific differential expression of exosomal miRNA in synovial fluid of patients with osteoarthritis. Sci Rep. 2017;7(1):2029.

60. Gao K, Zhu W, Li H, Ma D, Liu W, Yu W, et al. Association between cytokines and exosomes in synovial fluid of individuals with knee osteoarthritis. Mod Rheumatol. 2020;30(4):758–64.

61. Zhao Y, Xu J. Synovial fluid-derived exosomal lncRNA PCGEM1 as biomarker for the different stages of osteoarthritis. Int Orthop. 2018;42(12):2865–72.

62. Avci O, Unlu NL, Ozkumur AY, Unlu MS. Interferometric Reflectance Imaging Sensor (IRIS)--A Platform Technology for Multiplexed Diagnostics and Digital Detection. Sensors (Basel). 2015;15(7):17649–65.

63. Kowal J, Arras G, Colombo M, Jouve M, Morath JP, Primdal-Bengtson B, et al. Proteomic comparison defines novel markers to characterize heterogeneous populations of extracellular vesicle subtypes. Proc Natl Acad Sci U S A. 2016;113(8):E968–77.

64. Raposo G, Stoorvogel W. Extracellular vesicles: exosomes, microvesicles, and friends. J Cell Biol. 2013;200(4):373–83.

65. Andreu Z, Yanez-Mo M. Tetraspanins in extracellular vesicle formation and function. Front Immunol. 2014;5:442.

66. Caballero JN, Frenette G, Belleannee C, Sullivan R. CD9-positive microvesicles mediate the transfer of molecules to Bovine Spermatozoa during epididymal maturation. PLoS One. 2013;8(6):e65364.

67. Valadi H, Ekstrom K, Bossios A, Sjostrand M, Lee JJ, Lotvall JO. Exosome-mediated transfer of mRNAs and microRNAs is a novel mechanism of genetic exchange between cells. Nat Cell Biol. 2007;9(6):654–9.

68. Shao H, Im H, Castro CM, Breakefield X, Weissleder R, Lee H. New Technologies for Analysis of Extracellular Vesicles. Chem Rev. 2018;118(4):1917–50.

69. Turchinovich A, Weiz L, Langheinz A, Burwinkel B. Characterization of extracellular circulating microRNA. Nucleic Acids Res. 2011;39(16):7223–33.

70. Ali SA, Gandhi R, Potla P, Keshavarzi S, Espin-Garcia O, Shestopaloff K, et al. Sequencing identifies a distinct signature of circulating microRNAs in early radiographic knee osteoarthritis. Osteoarthritis Cartilage. 2020;28(11):1471–81.

71. Beyer C, Zampetaki A, Lin NY, Kleyer A, Perricone C, Iagnocco A, et al. Signature of circulating microRNAs in osteoarthritis. Ann Rheum Dis. 2015;74(3):e18.

72. Ntoumou E, Tzetis M, Braoudaki M, Lambrou G, Poulou M, Malizos K, et al. Serum microRNA array analysis identifies miR-140-3p, miR-33b-3p and miR-671-3p as potential osteoarthritis biomarkers involved in metabolic processes. Clin Epigenetics. 2017;9:127.

73. Rousseau JC, Millet M, Croset M, Sornay-Rendu E, Borel O, Chapurlat R. Association of circulating microRNAs with prevalent and incident knee osteoarthritis in women: the OFELY study. Arthritis Res Ther. 2020;22(1):2.

74. Zhang X, Wang C, Zhao J, Xu J, Geng Y, Dai L, et al. miR-146a facilitates osteoarthritis by regulating cartilage homeostasis via targeting Camk2d and Ppp3r2. Cell Death Dis. 2017;8(4):e2734.

75. Yamasaki K, Nakasa T, Miyaki S, Ishikawa M, Deie M, Adachi N, et al. Expression of MicroRNA-146a in osteoarthritis cartilage. Arthritis Rheum. 2009;60(4):1035–41.

76. Kang L, Yang C, Song Y, Liu W, Wang K, Li S, et al. MicroRNA-23a-3p promotes the development of osteoarthritis by directly targeting SMAD3 in chondrocytes. Biochem Biophys Res Commun. 2016;478(1):467–73.

77. Aae TF, Karlsen TA, Haugen IK, Risberg MA, Lian ØB, Brinchmann JE. Evaluating plasma extracellular vesicle microRNAs as possible biomarkers for osteoarthritis. Osteoarthritis and Cartilage Open. 2020;1(3-4):100018.

78. Seol D, McCabe DJ, Choe H, Zheng H, Yu Y, Jang K, et al. Chondrogenic progenitor cells respond to cartilage injury. Arthritis Rheum. 2012;64(11):3626–37.

79. Li Z, Huang Z, Bai L. Cell Interplay in Osteoarthritis. Front Cell Dev Biol. 2021;9:720477.

80. Muller M, Fazi F, Ciaudo C. Argonaute Proteins: From Structure to Function in Development and Pathological Cell Fate Determination. Front Cell Dev Biol. 2019;7:360.

81. Liu P, Zhang X, Li Z, Wei L, Peng Q, Liu C, et al. A significant role of transcription factors E2F in inflammation and tumorigenesis of nasopharyngeal carcinoma. Biochem Biophys Res Commun. 2020;524(4):816–24.

82. Pellicelli M, Picard C, Wang D, Lavigne P, Moreau A. E2F1 and TFDP1 Regulate PITX1 Expression in Normal and Osteoarthritic Articular Chondrocytes. PLoS One. 2016;11(11):e0165951.

83. Liu Y, Zou R, Wang Z, Wen C, Zhang F, Lin F. Exosomal KLF3-AS1 from hMSCs promoted cartilage repair and chondrocyte proliferation in osteoarthritis. Biochem J. 2018;475(22):3629–38.

84. van den Akker GGH, Caron MMJ, Peffers MJ, Welting TJM. Ribosome dysfunction in osteoarthritis. Curr Opin Rheumatol. 2022;34(1):61–7.

85. Chabronova A, van den Akker GGH, Meekels-Steinbusch MMF, Friedrich F, Cremers A, Surtel DAM, et al. Uncovering pathways regulating chondrogenic differentiation of CHH fibroblasts. Noncoding RNA Res. 2021;6(4):211–24.

86. Balaskas P, Green JA, Haqqi TM, Dyer P, Kharaz YA, Fang Y, et al. Small Non-Coding RNAome of Ageing Chondrocytes. Int J Mol Sci. 2020;21(16).

87. Peffers M, Liu X, Clegg P. Transcriptomic signatures in cartilage ageing. Arthritis Res Ther. 2013;15(4):R98.

88. Pimlott Z, Hontoir F, Ashraf Kharaz Y, Anderson J, Dyer P, Collins J, et al. Small nucleolar RNAs as mediators of oxidative stress in cross species cartilage and osteoarthritis. Osteoarthritis and Cartilage. 2020;28(1):S342.

89. Langhendries JL, Nicolas E, Doumont G, Goldman S, Lafontaine DL. The human box C/D snoRNAs U3 and U8 are required for pre-rRNA processing and tumorigenesis. Oncotarget. 2016;7(37):59519–34.

90. Peffers MJ, Fang Y, Welting TM, Haldenby S, Liu X, James V. Small RNA Sequencing reveals small non-coding RNA communication between synoviocytes and chondrocytes. United Kingdom Extracellular Vesicle Society; London 2019.

