## Supplementary material for "Small non-coding RNA landscape of extracellular vesicles from a post-traumatic model of equine osteoarthritis": Supplemantary File 1

Supplementary File 1. Plasma-derived extracellular vesicle size and size distribution. (A) Mode, (B) D10, (C) D50 and (D) D90 size of extracellular vesicles isolated from plasma at intervals between 0-63 days following model induction. All analyses were conducted via nanoparticle tracking using a Nanosight NS300. Error bars ± 1 standard deviation.

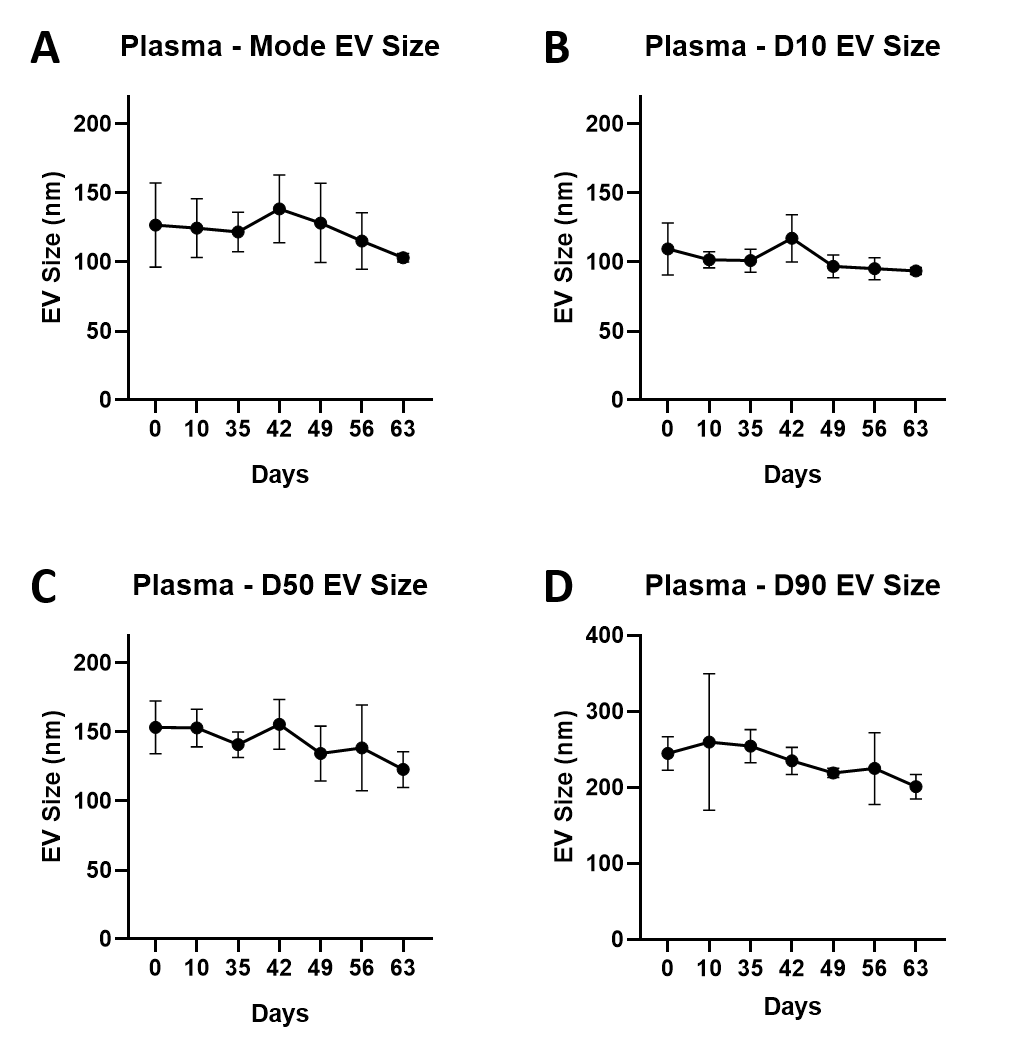
