## Supplementary material for "Small non-coding RNA landscape of extracellular vesicles from a post-traumatic model of equine osteoarthritis": Supplemantary File 2

Supplementary File 1. Synovial fluid-derived extracellular vesicle size and size distribution. (A) Mode, (B) D10, (C) D50 and (D) D90 size of extracellular vesicles isolated from synovial fluid control (green) and osteoarthritic (red) middle carpal joints at intervals between 0-63 days following model induction. All analyses were conducted via nanoparticle tracking using a Nanosight NS300. Error bars ± 1 standard deviation.

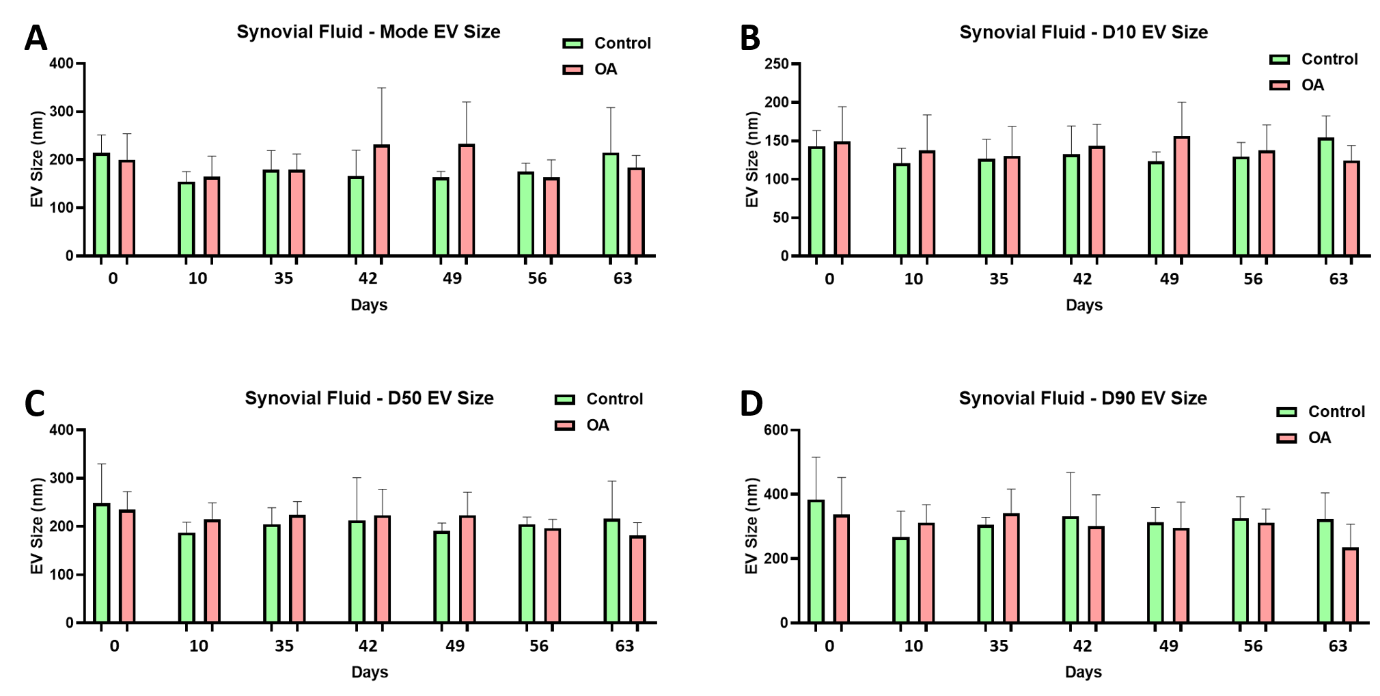
