## Supplementary material for "Small non-coding RNA landscape of extracellular vesicles from a post-traumatic model of equine osteoarthritis": Supplemantary File 3

Supplementary File 3. Plasma derived EV colocalisation results for tetraspanins CD9 and CD81. A) Day 0, B) Day 42, C) Day 63. Fourescent channels for CD81 (red) and CD9 (blue) are shown. The ratio of CD81/CD9 is in yellow.
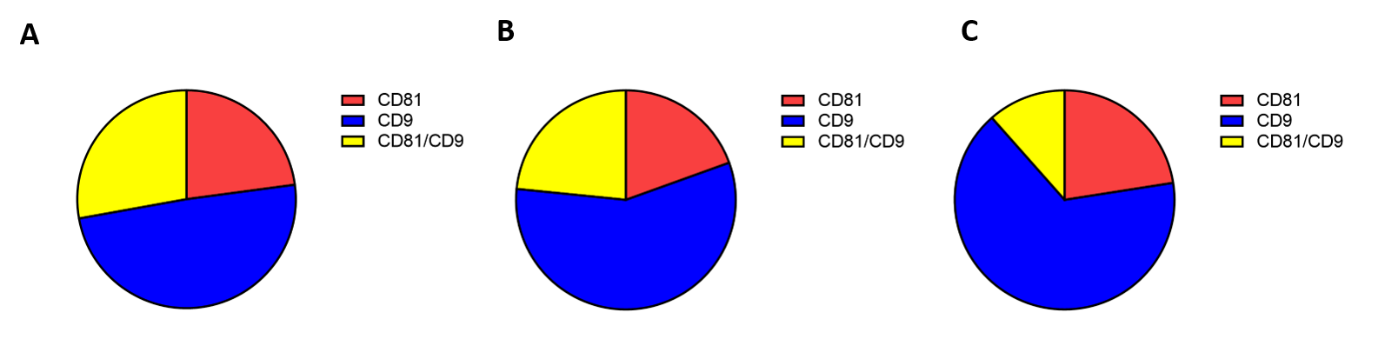
