## Supplementary material for "Small non-coding RNA landscape of extracellular vesicles from a post-traumatic model of equine osteoarthritis": Supplemantary File 4

Supplementary File 4. Classes of non-coding RNAs identified in groups in which they were found in at least 30% of all samples within that group.

| **Sample** | **Type** | **Number** |
| --- | --- | --- |
| Plasma | lncRNA | 47 |
| Plasma | miRNA | 52 |
| Plasma | snoRNA | 6 |
| Plasma | snRNA | 17 |
| Plasma | tRNA | 36 |
| Synovial Fluid Control | lncRNA | 66 |
| Synovial Fluid Control | miRNA | 77 |
| Synovial Fluid Control | snoRNA | 4 |
| Synovial Fluid Control | snRNA | 18 |
| Synovial Fluid Control | tRNA | 51 |
| Synovial Fluid OA | lncRNA | 85 |
| Synovial Fluid OA | miRNA | 74 |
| Synovial Fluid OA | snoRNA | 4 |
| Synovial Fluid OA | snRNA | 20 |
| Synovial Fluid OA | tRNA | 50 |
