## Supplementary material for "Small non-coding RNA landscape of extracellular vesicles from a post-traumatic model of equine osteoarthritis": Supplemantary File 5

Supplementary File 5. Differentially expressed miRNAs isolated from synovial fluid-derived extracellular vesicles. Error bars ± 1 standard deviation. ∗ = p < 0.05, ∗∗ = p < 0.01.

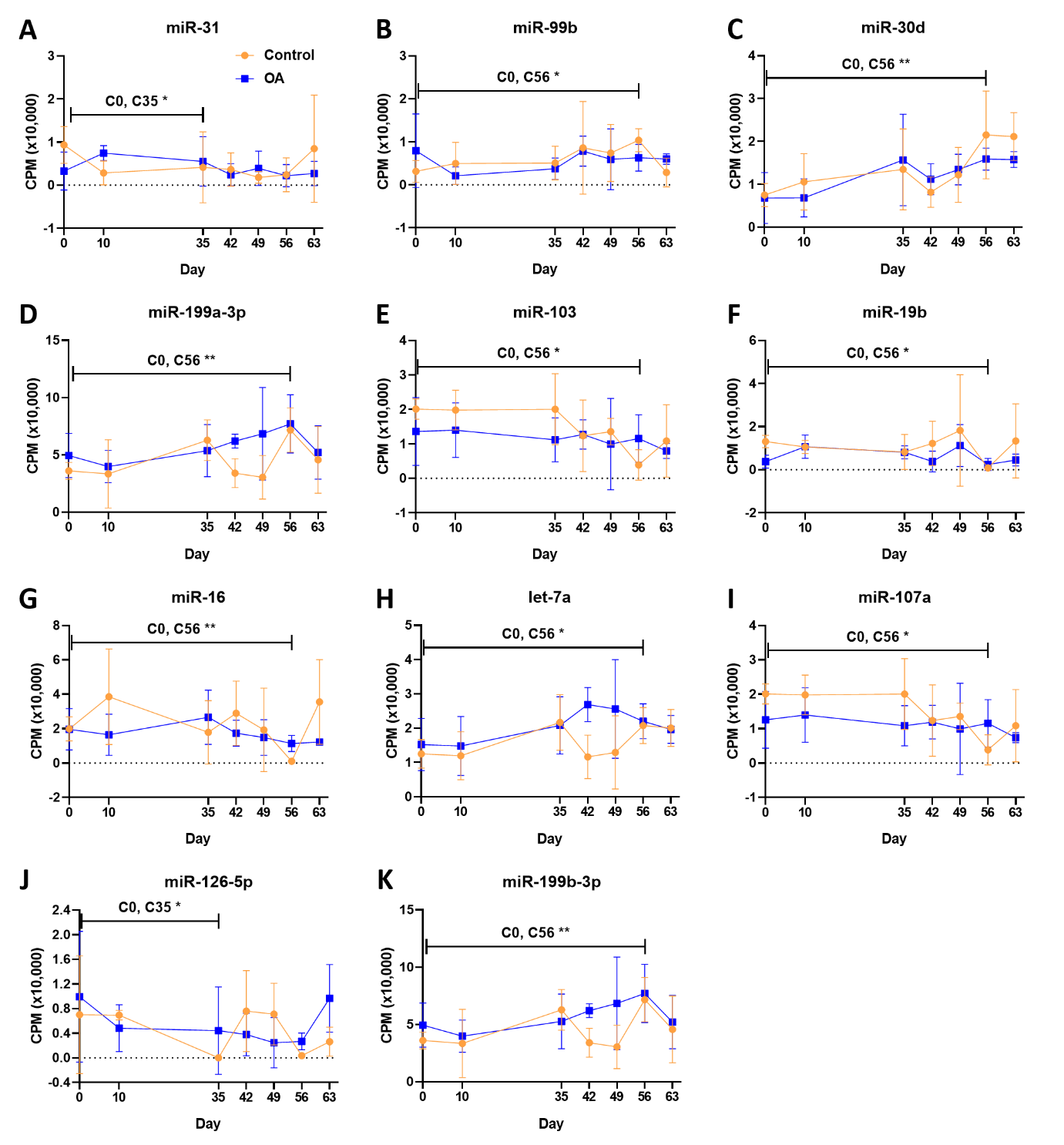
